# Premotor cortex uses a compositional neural geometry to plan words

**DOI:** 10.64898/2026.04.27.721195

**Authors:** Benyamin Abramovich Krasa, Erin M. Kunz, Foram Kamdar, Donald Avansino, Nick Hahn, Akansha Singh, Nicholas S. Card, Maitreyee Wairagkar, Carrina Iacobacci, Leigh R. Hochberg, David M. Brandman, Sergey D. Stavisky, Jaimie M. Henderson, Francis R. Willett, Shaul Druckmann

## Abstract

Speech requires precise serial ordering of words and phonemes into novel combinations. To accomplish this, the brain is believed to flexibly prepare utterances before producing them, even allowing pronunciation of never-before spoken words. To discover how neural populations achieve this, intracortical activity from premotor cortex was recorded while two speech neuroprosthesis pilot clinical trial participants attempted to speak factorially-balanced phoneme sequences. During preparation, activity encoded not only the next-phoneme, but multiple upcoming phoneme positions spanning whole words. We found that word-level plans were formed by compositionally combining phoneme representations, a mechanism that may enable efficient planning of novel sequences. When utterances contained more than one word, premotor cortex activity was largely limited to the first word, suggesting that articulatory planning is segmented by higher-order features. Together, these results reveal a compositional, hierarchically-segemented planning geometry, potentially a universal neural strategy for sequence organization across higher levels of language.

## Introduction

Speech is generated by the complex control of articulation, requiring coordination of multiple effectors (e.g., lips, tongue, larynx, diaphragm) to produce precise sequences of sound. The motor plan underlying the sequencing of articulatory gestures is hypothesized to be stored in neural activity as an “articulatory buffer”^1–4^. Support for this theory comes from both behavioral studies of phonological errors in natural speech^5^ (e.g. spoonerisms such as erroneously saying *blushing crow* instead of *crushing blow*), case studies of deficits in articulatory sequencing due to stroke^6,7^, neural recordings^8–10^, and modeling^11,12^. These studies indicate that speech is hierarchically planned and have proposed anatomical hypotheses for where motor programs of speech sequences are represented, but have so far lacked the resolution to directly test the underlying representation of an articulatory buffer in neural population activity.

In nonhuman primate and rodent animal models, the neural population mechanisms underlying motor preparation have been studied extensively^13,14^ (e.g., for behaviors such as reaching or licking) and provide hypotheses for speech planning. These studies have found that persistent neural activity during an instructed delay period contains a low-dimensional representation of the upcoming movements, which has been interpreted as a motor plan(Fig. 1A). This preparatory activity is predictive of single trial kinematics and reaction times, and has been causally linked to the upcoming movement^15–18^. One study of reaching sequences found that preparatory activity in motor cortex was limited to the first planned movement, and that subsequent movements were independently planned and executed after a go cue^19^. Alternatively, a study of eye-saccade sequences found that in the lateral prefrontal cortex, whole sequences of movements were simultaneously prepared using orthogonal subspaces for each sequence position and larger representations for more immediately upcoming positions^20^. How this framework might extend to ventral premotor areas during articulatory or word sequence planning in humans remains unknown.

**Figure 1:**
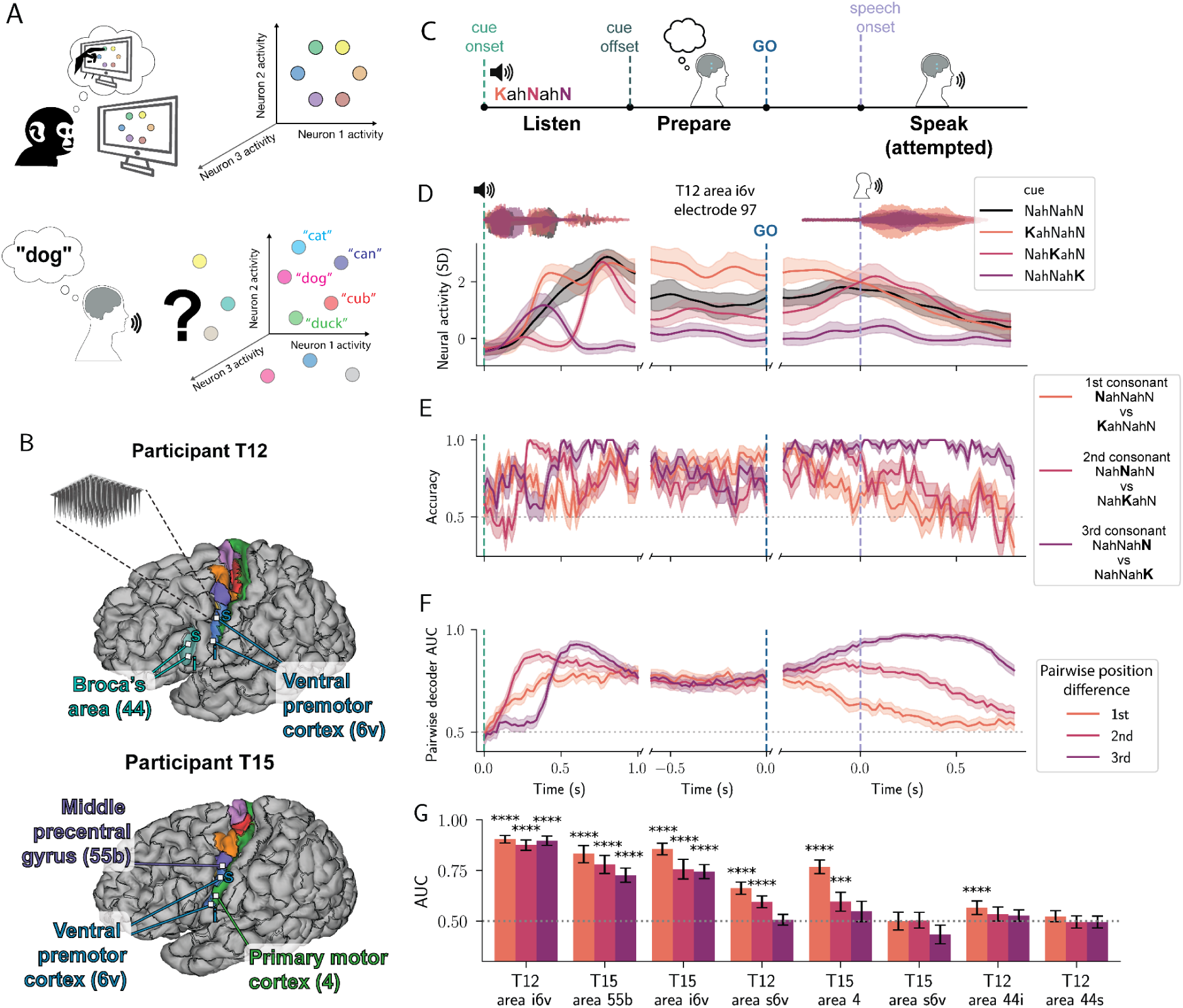
Preparatory activity for speech encodes whole sequences of articulatory gestures. **A)** Schematic of previously characterized neural geometry of motor preparation in primate reaching (top) and hypothesized geometry in speech preparation (bottom). **B)** Microelectrode array locations (white squares) with HCP-identified areas highlighted on the cortical surface. **C)** Schematic of instructed-delay task structure. Sequences of alternating consonants and vowels were cued to participants via audio alone during the listening epoch of a trial. **D)** Smoothed neural activity (threshold-crossing counts) recorded on a single example electrode in T12 area i6v in response to three phoneme sequences that differ in either the 1st (orange), 2nd (red), or 3rd (purple) sequence position from an example base NahNahN condition (black). Trial epochs were aligned separately in each of the axes to: audio cue onset (left), preparatory window before the go cue after audio cue offset (middle), and speech onset (right). Shaded intervals are 95% CI of the mean bootstrapped over trials. Microphone audio traces of the cue and attempted speech envelopes are plotted above. **E)** Decoding accuracy over time of LDA decoders fit on i6v data to decode between pairs of conditions from **D** that differed in only one position. Average accuracy across held out trials is plotted. Shaded intervals are standard error of the mean over held out trials **F)** Decoding performance (Area Under the Receiver Operating Characteristic Curve (AUC)) over time of LDA decoders fit to all pairs of conditions that differ in only one position, averaged across all pairs. Shaded intervals are bootstrapped 95% CI of the mean over all pairs. **G)** Summary of LDA decoder performance for all pairs that differed in one position based on a 0.8s window of preparatory activity for each brain region across participants. Error bars are bootstrapped 95% CI over decoders. \*\*\**P*<0.001, \*\*\*\**P*<0.0001, one-sample, one-sided t-test with FDR correction over decoders.

Recent work on the neurophysiology of speech has highlighted sustained activity in areas of premotor cortex during speech preparation, but these studies have not yet investigated the content and geometry of these preparatory representations at a neural population level with cellular resolution^8–10^. To do so would require experimental paradigms that systematically span a large enough number of phoneme combinations, combined with neural recordings that can measure population activity at single neuron resolution. In our previous work as part of the BrainGate2 pilot clinical trial, we have shown that intracortical microelectrode recordings from premotor cortex in people with paralysis during attempted speech can be successfully decoded into text, thus restoring communication^21–24^. At the same time, the underlying structure of neural activity encoding phoneme sequences remains unknown, and its characterization could inform future speech brain-compute interface (BCI) design.

Here, we recorded neural activity from two of these same participants during full-factorial instructed-delay tasks that systemically varied the identity of phonemes in each serial position of a word. We report persistent preparatory activity before speech onset that encodes whole sequences of phonemes using a compositional code, with a planning horizon that is largely limited to the first word in a sequence. We also demonstrate that the neural representation of phonemes during audio cue presentation, preparation, and attempted speech is correlated, in contrast to prior results found in macaques for reaching movements. These results are evidence for the existence of an articulatory buffer in precentral gyrus and provide a first characterization of its neural geometry, revealing distinct preparatory motifs from those described in animal models, and suggesting a general mechanism for flexible sequence planning across the hierarchy of language planning.

## Results

### Preparatory activity encodes whole sequences of articulatory gestures

To investigate the role of preparatory activity underlying speech, neural activity was recorded from microelectrode arrays in two participants while they prepared and attempted to speak instructed utterances. Activity was recorded from 64-channel microelectrode arrays (Blackrock Neurotech, Salt Lake City, UT, USA) in speech areas of precentral gyrus in two participants (T12 and T15, Fig. 1B) enrolled in the BrainGate2 clinical trial. Both T12 and T15 were dysarthric at the time of this study due to amyotrophic lateral sclerosis (ALS). Arrays were targeted using the Human Connectome Project (HCP) parcellation procedure^25^ applied to presurgical structural and functional resting-state MRI data. Arrays were placed in the inferior and superior region of HCP-identified area 6v (referred to as i6v and s6v, respectively, in premotor cortex) as well as the inferior and superior region of HCP-identified area 44 (putative Broca’s area) in T12. For participant T15, arrays were placed in HCP-identified areas i6v, s6v, 4 (primary motor cortex), and 55b (middle precentral gyrus). See Figure 1B for anatomical locations and HCP areal boundaries.

We focused on phonemes as an instructable unit of articulatory gestures. To systematically study the effects of phoneme identity and sequence position, we designed a full-factorial articulatory sequence paradigm. In an instructed-delay task, participants listened to nonsense words composed of sequences of alternating consonants and vowels (CVCVC), with a limited set of articulatorily distant consonants (e.g. /K/, /N/, /SH/, /T/; ARPABET phoneme notation) and a single vowel (/AH/). Nonsense words were used so that all possible consonant combinations of a given length could be tested in a full factorial design (Fig. S1). After a delay period, an audio-visual go cue instructed participants to attempt to speak and reproduce the cued audio (Fig. 1C). Two estimates of local neuronal action potential activity were computed for each electrode (threshold-crossing counts and spike-band power, see Methods for details). In T12, activity on 186 out of 256 electrodes significantly differed from rest during at least one condition (Welch’s t-test with False Discovery Rate (FDR) correction, α=0.05; T15: 217/256 electrodes).

To assess how neural activity represented each sequence position independently, we compared pairs of sequences that differed only in a single position (e.g., the pair **K**ahNahSH vs **N**ahNahSH, which differ only in the 1st position). Sequence positions (1st, 2nd, 3rd) refer to consonant position (C_1_VC_2_VC_3_) as vowels did not differ between conditions, though it should be noted that the full sequence contains five phonemes so the operationalized 3rd sequence position here actually refers to the fifth phoneme. In line with nonhuman primate studies of preparatory activity underlying arm reaches^19,26,27^, neural responses of single electrodes during the preparatory period were significantly different between pairs of sequences that differed in the immediately upcoming movement, i.e. the 1st consonant (T12: 103/256 electrodes, T15: 54/256; Welch’s t-test with FDR correction, α=0.05). More surprisingly, and unlike some previous findings for arm movement sequences, many single electrodes had significantly different activity during the delay period for sequences that differed only in the 2nd or 3rd position while being identical in the 1st position (example electrode shown in Figure 1D; 2nd position: T12: 131/256, T15: 99/256; 3rd position T12: 83/256, T15: 89/256; Welch’s t-test with FDR correction, α=0.05).

To assess sequence encoding at the neural population level, linear discriminant analysis (LDA) decoders were fit to distinguish pairs of sequences that differed in the 1st, 2nd, or 3rd sequence position (Fig. 1E, see Methods). All three positions could be decoded significantly above chance from areas i6v and 55b during a preparatory period before the go-cue (Fig. 1G; T12 area i6v *P*=[4.99e-75, 3.04e-62]; T15 area 55b *P*=[3.61e-18, 3.00e-08]; T15 area i6v *P*=[3.78e-22, 2.02e-9]; one-sample t-test over decoders, FDR corrected, range of *P* values for all three sequence positions reported). Other nearby regions had above chance decoding only for the 1st and 2nd consonants (T12 area s6v, T15 area 4; Fig. 1G). In summary, we found that neural preparatory activity for speech in areas i6v and 55b simultaneously encoded sequences of articulatory plans for whole pseudo-words prior to execution of the first consonant, rather than just encoding the very first articulatory gesture prior to movement (and only later during production encoding subsequent elements). In contrast, other nearby speech-related regions’ tuning mostly (but not solely) encoded the initial articulation (Fig. 1G, T12 area s6v, T15 area 4).

### A correlated representation of phonemes across sequence positions

The simultaneous encoding of all three consonant positions raises the question: what is the neural structure underlying the articulatory plan for whole words? Previous studies in lateral prefrontal cortex found a neural population state geometry whereby each sequence position is encoded in a separate subspace and these subspaces are orthogonal across different sequence positions^20^. In contrast, analyzing activity from T12 area i6v during a 0.7s preparatory window (i.e., before go cue), we found that nearby sequence positions were not always orthogonally represented. For example, decoders trained only on 3rd sequence position identity were able to predict 2nd position differences (AUC = 0.73, 95% CI [0.71, 0.74], *P*<0.0001, permutation test), and vice versa (AUC = 0.75, 95% CI [0.73, 0.77], *P*<0.0001, permutation test; Fig. 2A). Similar generalization was observed across multiple regions (Fig. S2). Together, these findings demonstrate that preparatory phoneme representations are not organized into orthogonal subspaces across positions.

**Figure 2:**
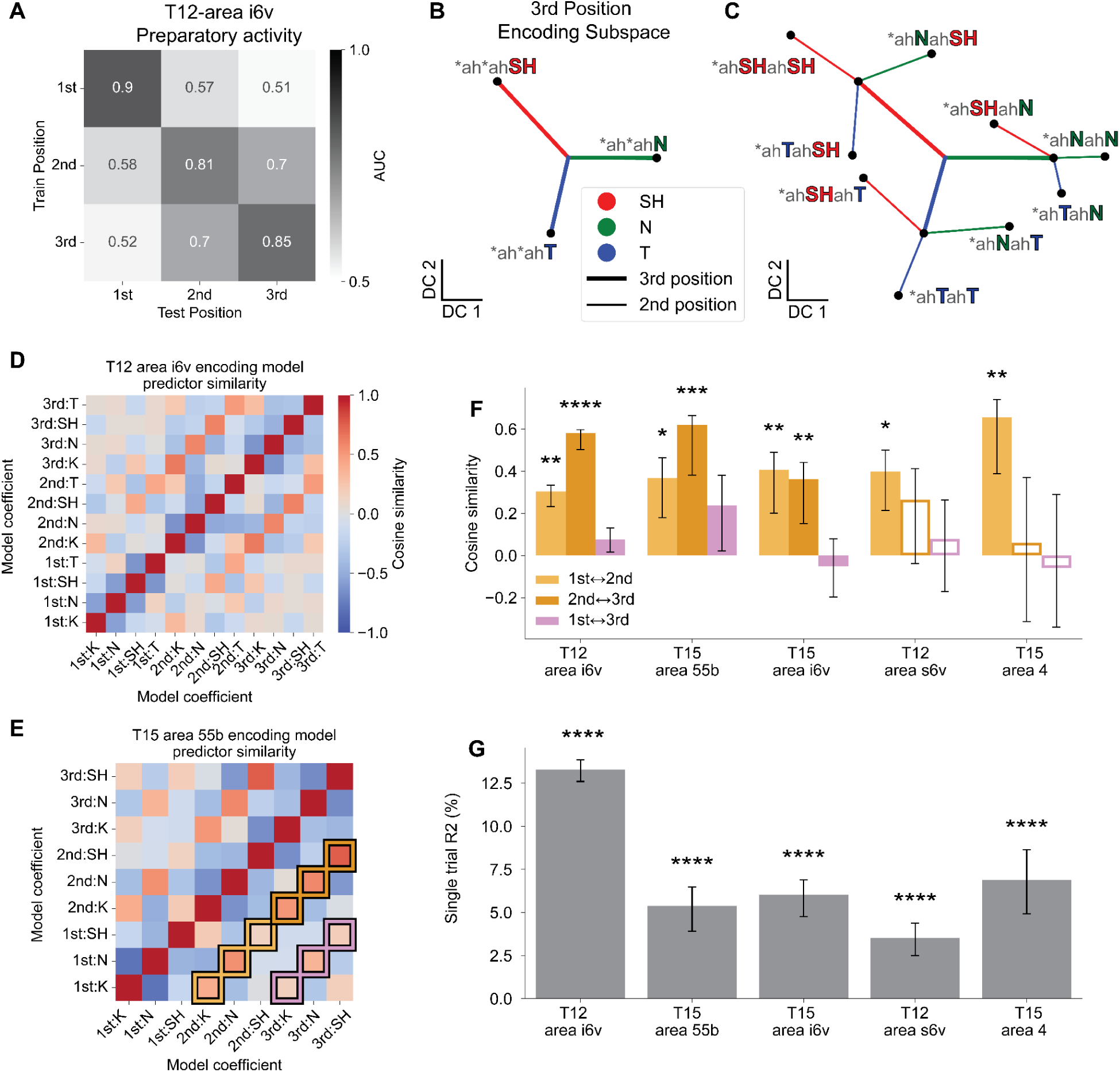
Preparatory representations of phonemes are correlated across positions. **A)** Multiclass LDA decoders were fit to predict the phoneme in each sequence position and tested in all other sequence positions to assess representational similarity across positions (0.7s preparatory window; T12 area i6v). **B)** Average representation of 3rd position phonemes in held out trials projected into discriminant component 1 (DC1) and DC2 describes an encoding space for 3 class decoding between sequences ending in SH, N, or T. *\*ah*ah*SH* refers to all sequences ending in /SH/ (e.g. *KahNahSH*, *SHahNahSH*, etc) **C)** Trial-averaged projection for pairs of 2nd and 3rd consonants (marginalized over 1st-position), for the same encoding space in B. Lines are displacement-vectors to visualize encoding geometry for 3rd position (thick lines) and 2nd position independent of 3rd position (thin lines). See Figure S3 for explanation of analysis. **D)** Pairwise cosine similarity between coefficients of a linear encoding model to predict preparatory neural activity in area i6v in T12 as a linear sum of phoneme representation vectors in each position. **E)** Same as **D** for T15 area 55b. Overlaid boxes on lower off-diagonal highlight pairs of like-phonemes across positions summarized in **F**. **F)** Correlated representation across positions estimated as cosine similarity. Pairwise cosine similarity for like-phoneme-model coefficients were grouped by position pair (e.g. 1st ↔ 2nd) and averaged. All arrays tuned to at least two positions were included. Unfilled bars indicate one of the positions in the comparison was not decodable above chance in Figure 1G. **G**) Linear encoding model performance across arrays. Single-trial R^2^ is the variance explained for model predictions of electrode activity for single trials of held-out data. Error bars in **F** and **G** are bootstrap 95% confidence intervals, statistical significance assessed via permutation test. Significance is denoted by asterisks: \**P*<0.05, \*\**P*<0.01, \*\*\**P*<0.001, \*\*\*\**P*<0.0001

Population activity typically cannot be visualized in its original, high dimensional state space and thus targeted dimensionality reduction is used to visualize the structure^28–30^. Linear discriminant analysis (LDA) finds an encoding subspace of a maximal dimension equal to the number of classes minus one^31^. As we have more than four conditions, the neural encoding of whole sequences could not be directly visualizable if we fit an LDA model on the full set of conditions. However, by instead fitting an LDA model to a single position, i.e., discriminating between 3 phonemes in that position, we obtain a two dimensional subspace and can directly visualize the projected neural data. We start with the two dimensional subspace defined by an LDA model fit to decode the 3rd position, which captures 13.6% of the condition-averaged variance in held-out data. Projecting neural activity from held-out trials into this subspace and analyzing the contribution of different positions to the neural geometry, we found a generalized representation of consonants across sequence position (Fig. 2BC, see Fig. S3 and Methods for additional visual explanation of the analyses). First, as expected from the high decoding performance for the 3rd position, trials grouped and averaged according to the 3rd position consonant were well separated in the decoding space (Fig. 2B, see Fig. S3E for single trials). Next, to visualize the relationship between 2nd and 3rd position encoding geometry, 2nd position displacement vectors were estimated. These displacement vectors represent 2^nd^ position consonant identity as directions in neural state space, estimated relative to the mean activity for each 3rd position consonant — providing a geometric test of whether 2nd position identity is encoded consistently across sequence positions. Rather than only grouping trials by the identity of the 3rd position, trials were grouped by 2nd and 3rd position sequences (e.g. breaking *ah*ahSH into the component, *ahNahSH, *ahSHahSH, *ahTahSH; asterisk indicates marginalizing over consonants in other sequence positions; see Fig. S3D for methodological details). The displacement due to the 3rd position was similarly oriented as the 2nd position (Fig. 2C, compare thick and thin lines of the same color). That is, the 9 task conditions (3 positions x 3 consonants) formed a compositional structure, with neural population activity vectors encoding the 2nd position largely independent of the specific 3rd position consonant and closely aligned with vectors encoding the same phoneme when it was in the 3rd position (Fig. 2C, single trials shown in Fig. S3EF). Notably, this geometry emerges from only fitting to the 3rd position, demonstrating a consistent, correlated representation across sequence positions and not an orthogonal subspace for each position.

The visualization of the geometry suggests a position-invariant linear encoding model, where the pattern of activity corresponding to a particular sequence would be well described by an encoding largely independent of the full context of the sequence, i.e., by a linear sum of vectors each independently describing the contribution of an individual phoneme at a given position. Formally, the predicted activity (*y*) for a sequence with phonemes (*p*_1_, *p*_2_, *p*_3_) is a sum of position-specific weight vectors (β): *y* = *p*_1_ β_1_ + *p*_2_ β_2_ + *p*_3_ β_3_ (see Methods). We found that such linear models fit to predict activity on significantly tuned electrodes accounted for a substantial fraction of neural variance, explaining 56.8% of condition-averaged variance in held-out data from T12 area i6v (conditions defined by 2nd-3rd position consonant identity i.e. marginalizing over 1st position; 13.3% for held-out single trials; *P*<0.0001, permutation test). These values indicate that a simple linear composition of phoneme and position representations captures a meaningful component of preparatory population activity. The compositional model is a substantial reduction in complexity from uniquely representing each sequence (e.g. for 4 phonemes in 3 positions, the required set of encoding vectors is reduced from 4^3^ = 64 to 4×3=12). Consistent with the visualization in Fig. 2C, we find that in T12 area i6v, coefficients for the same phoneme across adjacent positions were significantly correlated (Fig. 2D), with the 2nd and 3rd positions better aligned than the 1st and 2nd (cosine similarity 0.58 [0.50, 0.60] versus 0.30 [0.23, 0.33] for 2nd↔3rd and 1st↔2nd respectively; see Figure S4 for visualization). A similar trend was observed in T15 area 55b, with weaker correlations between the 1st and 2nd position than between the 2nd and 3rd positions (Fig. 2E; cosine similarity 0.62 [0.38, 0.66] versus 0.37 [0.18, 0.46], see Fig. S2C). This suggests the 1st position is encoded in a more separate subspace, likely to facilitate its use for driving the upcoming articulatory gesture. The same linear model was fit to all arrays that contained significantly decodable representations across multiple positions to assess whether this phenomenon generalizes across speech-motor areas. Consistently, in all arrays tuned to multiple positions, phonemes in adjacent positions were similarly represented (Fig. 2FG. cosine similarity between adjacent position coefficients: T15 i6v: 1st↔2nd: 0.41 [0.20, 0.49], *P*=0.0028; 2nd↔3rd: 0.36 [0.15, 0.44], *P*=0.0075; T12 s6v: 1st↔2nd: 0.40 [0.20, 0.49], *P*=0.012; T15 area 4: 1st↔2nd: 0.66 [0.39, 0.74], *P*=0.0012, permutation test), whereas non-adjacent position pairs (1st↔3rd) were not significant in any area (all *P*>0.09). Across all areas, the compositional linear model captured a significant fraction of single-trial neural variance (3.5%–13.3%, *P*<0.0001 for each area; Fig. 2G). We note that this linear encoding results in a geometry that would benefit from non-linear decoding, such as a two stage decoder that subtracts out the contribution of one position before decoding the other.

### Preparatory activity represents both sequence length and phonetic content

Spoken words vary widely in both phonemic content and utterance length. Psycholinguistic studies have shown that speech onset times are delayed for longer words, suggesting that motor planning must encompass the articulatory length of an upcoming word before speech can be initiated^32^. It remains unclear whether the regions that encode phoneme sequences also represent utterance length.

To probe whether sequence length is represented in motor cortical preparatory activity, we presented sequences of varying length (CV, CVC, CVCVC consonant sequences) in a similar instructed-delay trial design with cues constructed in a full factorial design from a subset of consonants (Fig. 3A). First, we asked whether preparatory activity across multiple cortical areas was tuned to sequence length by fitting LDA models to decode sequence length only. Areas i6v and s6v in T12, and areas i6v and 55b in T15, showed significant sequence length decoding (T12 i6v: LDA normalized balanced accuracy = 0.68, 95% CI [0.628–0.731]; T12 s6v: 0.220 [0.177–0.265]; T15 i6v: 0.247 [0.153–0.345]; T15 55b: 0.625 [0.552–0.692]; Chance=0; all *P*<0.0008, permutation test with Bonferroni correction; see Methods), demonstrating that preparatory activity encodes utterance length across multiple regions of precentral gyrus.

**Figure 3:**
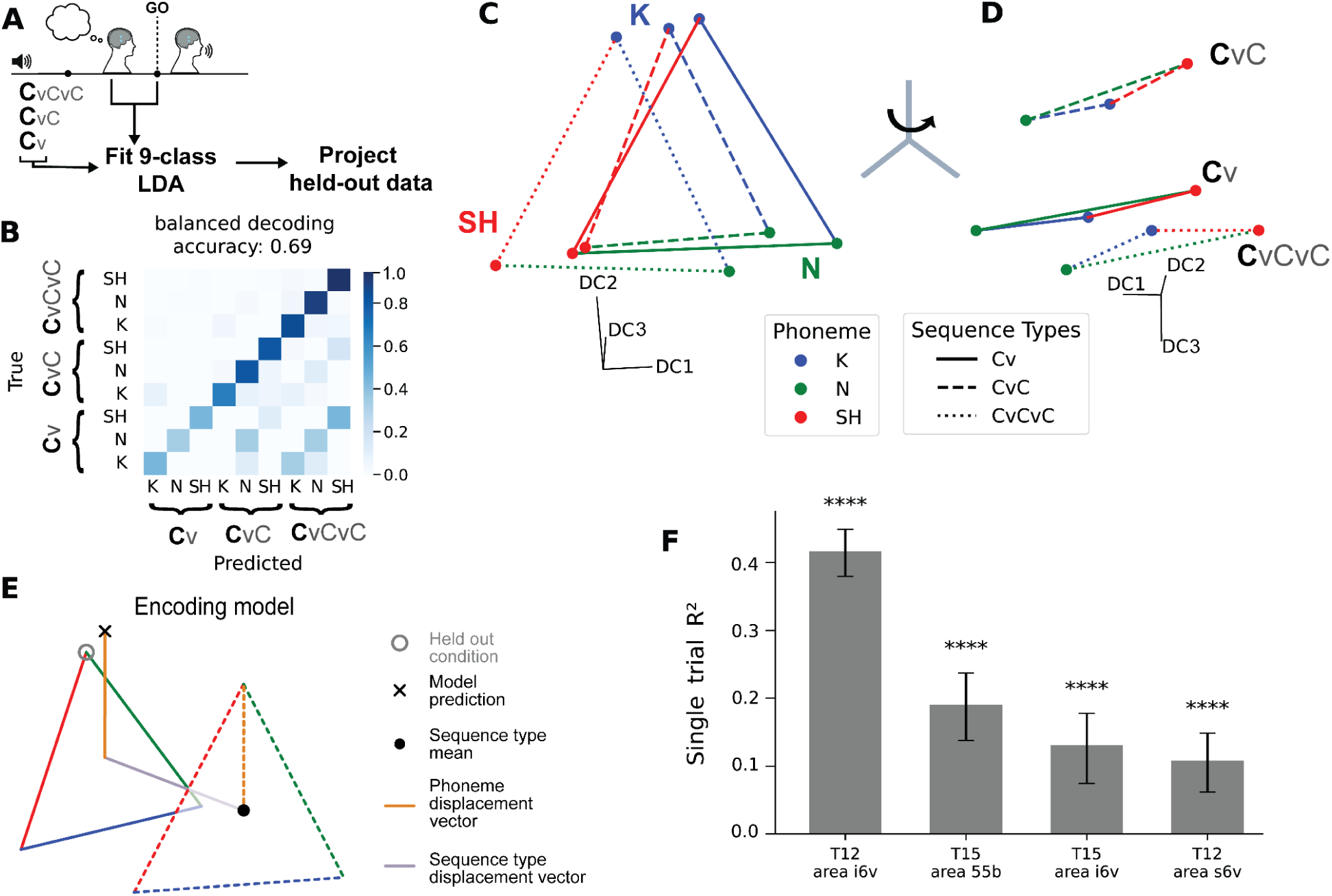
Both phonemic content and sequence length are simultaneously encoded in preparatory activity via a compositional geometry. **A)** In addition to CVCVC sequences, full factorial CVC and CV consonant sequences were cued with the same subset of consonants and fixed vowels in the same trial design. From a 0.6s window of preparatory activity in T12 area i6v, a multiclass LDA model was fit to predict 1st phoneme and sequence length from neural activity (threshold-crossing counts and spike-band power, 9-classes). **B)** Confusion matrix for 9-class prediction of 1st phoneme identity across three sequence lengths. Rows are normalized to sum to 1 due to class imbalance **C)** Projection of held out data into top three discriminant components (DCs) from the LDA model, averaged by class label. Lines connect like-sequences. Colors indicate the 1st phoneme in the sequence. Dimensions are rotated to highlight similar encoding of the phonemes across different sequence lengths. **D)** A rotation of the same data in **C** reveals a separate dimension that encodes sequence length. **E)** Schematic of a compositional encoding model to assess phoneme-representational similarity across sequence lengths. Model predictions of held-out classes were estimated from sequence-length and phoneme displacement vectors from all other conditions.**F)** Encoding model performance for all arrays. Single-trial R^2^ is the variance explained for model predictions of single trials projections in all DCs, held-out data. Error bars are bootstrap 95% confidence intervals, statistical significance assessed via permutation test. Significance is denoted by asterisks: \*\*\*\**P*<0.0001

Having established that sequence length is decodable across multiple areas, we next asked how it is represented alongside phoneme identity. First, we trained an LDA decoder to jointly classify both 1st-position phoneme identity and sequence-length across all nine combinations, achieving significantly above-chance performance (Fig. 3B; 69% balanced accuracy, 95% CI [62, 76], *P*<0.0001 permutation test). We note that in the factorial design with fixed trial count per sequence, shorter sequences had overall fewer trials; when controlling for this imbalance the confusion matrix was more uniform across sequence lengths (Fig. S5). To visualize the neural dimensions underlying the decoder’s performance, held-out trials were class-averaged and projected into the top three discriminant components of the LDA model (see Methods). Rather than uniform separation between classes, we observed a compositional arrangement of phoneme and sequence length. Sequences with the same 1st phoneme but different lengths were encoded similarly along one dimension, while a rotation within the three-dimensional space revealed a largely separate dimension encoding sequence length (Fig. 3CD).

We next asked whether the compositional geometry observed in Figure 3CD held for neural activity projected into the full discriminant-component space and generalized to other brain areas. To do so, we constructed a compositional prediction of held-out conditions by combining independently estimated phoneme and sequence-type components. Specifically, for a held-out sequence (e.g., a CVCVC sequence beginning with K), we estimated its neural representation as the sum of (i) a phoneme displacement vector derived from a different sequence length (e.g., a CVC sequence beginning with K) and (ii) a sequence-type displacement vector. Under a compositional code, phoneme-specific displacements should be preserved across sequence lengths, such that the predicted representation lies close to the true held-out activity. Consistent with this prediction, the compositional model closely matched observed neural representations. In T12 area i6v, the model accounted for 95.3% of the condition-averaged variance for neural activity projected into the full LDA model discriminant components space (95% CI [92.0, 96.3], *P*<0.0001; conditions defined by sequence-length and first consonant identity, marginalizing over 2nd and 3rd position elements; single-trial variance explained: 41.6% [38.0, 44.9] *P*<0.0001, permutation test). Additionally, models across all areas with preparatory activity tuned to sequence length was significantly above chance (single-trial variance explained: T12 s6v: 10.8% [6.2, 14.9]; T15 i6v: 13.1% [7.5, 17.8]; T15 55b: 19.1% [13.8, 23.7], all *P*<0.0001; permutation test). Beyond just the first sequence position, other sequence positions were also similarly represented across sequence types, when marginalizing out sequence-type representations (Figure S6). Together, these results indicate that preparatory activity in precentral gyrus encodes both phoneme sequences and utterance length, with length represented along partially separable dimensions that may complement overlapping phoneme representations across sequence positions to ensure reliable planning of sequences of the intended length.

### A generalized representation of phoneme sequences across audio-cue listening, motor preparation, and speech production epochs

Having characterized the structure of preparatory representations, we next asked how they relate to activity during perception and production. In terms of preparation and execution, prior work has suggested that preparatory activity can be prevented from spreading to downstream areas and prematurely triggering movement by constraining activity to a specific nullspace, orthogonal to that of execution activity^13,33–35^ (Fig. 4A). Under this hypothesis, decoders trained during preparation would not generalize to execution. In terms of perception and preparation, one might expect a reorganization of perception-time encoding into an articulation-driving plan, in particular as the full sequence is only revealed sequentially during perception and the whole sequence is represented in preparation. That is, the neural representation of, say, the third consonant may differ fundamentally between the moment it is first heard — when it is being parsed from an acoustic signal in the context of preceding sounds — and the moment of speech preparation, when it must be integrated into a coherent motor plan for the entire word. If these results hold true for speech preparatory activity, then a neural decoder fit to activity during different periods would not be able to generalize well to data during other periods (Fig. 4B).

**Figure 4:**
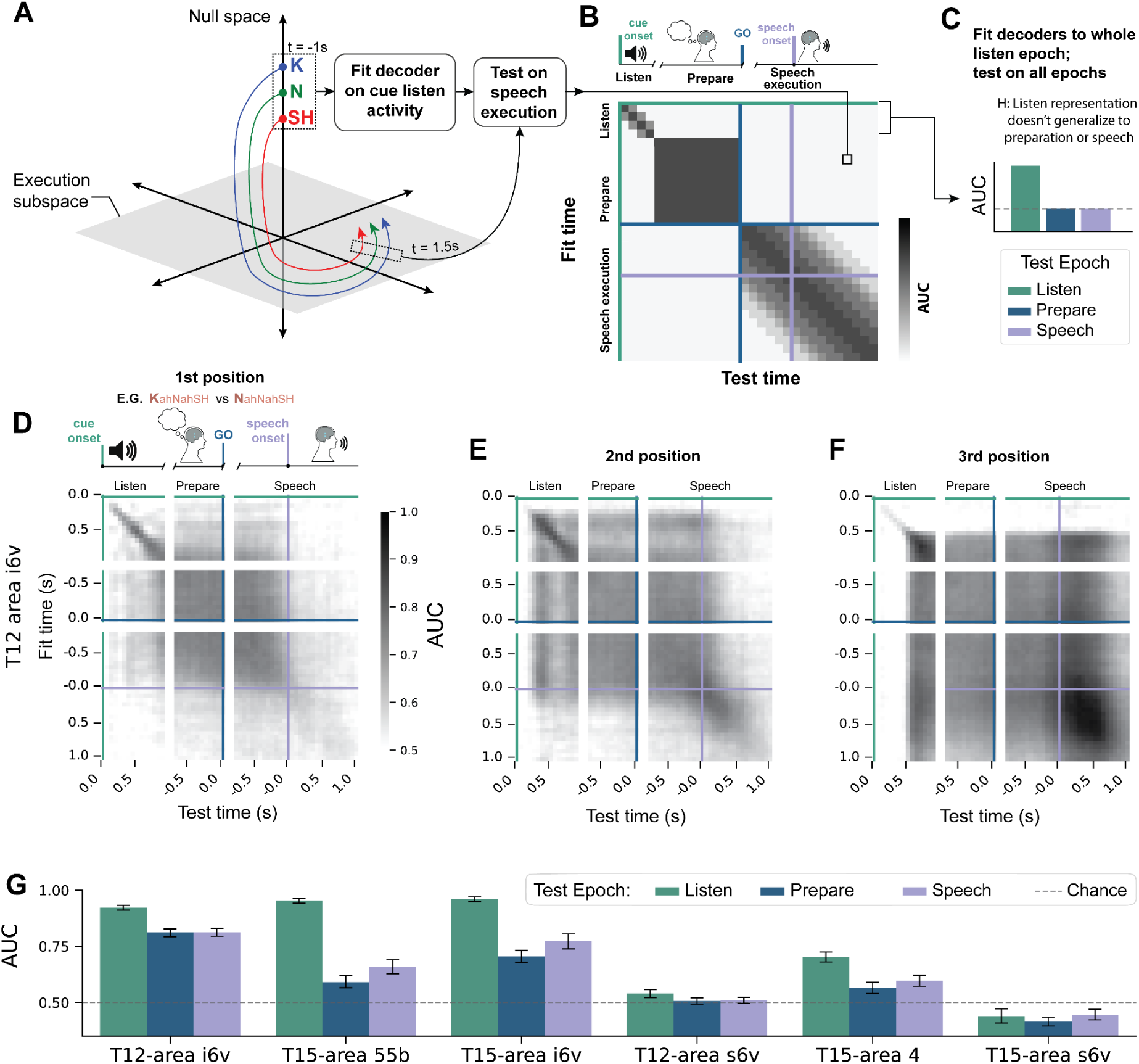
Phoneme sequence representation persists across trial epochs. **A)** Schematic of the neural geometry and trajectories of preparatory activity inspired from studies of arm reaching. Hypothesized cue-listening activity for different phonemes is organized along a null-space neural dimension that lies orthogonal to an execution subspace. **B)** Schematic of orthogonal hypothesis prediction: a linear decoder fit to audio-cue listening activity does not generalize to neural activity during attempted speech. The diagram as a whole represents one hypothesis for decoder generalization analysis results for **D, E, F** where neural representations rapidly change as audio cue is heard and interpreted, a stable preparatory representation emerges after audio cue onset, and an orthogonal and continually changing neural representations emerges during speech **C)** A schematic summary of hypothesized decoder generalization matrices. Decoders fit during the audio cue listening epoch are predicted to have high performance on held out trials during the listen epoch (same as fit context), but perform poorly when assessed on preparatory or attempted speech epochs due to orthogonality between representations. Bar plot is a hypothesis for **G**. **D)** Decoder generalization matrix fit on neural activity from T12 area i6v. Decoders were fit on 100ms windows (60ms slide) of neural activity to discriminate between condition pairs that differ in only the 1st sequence position. Average AUC over pairs is shown **EF)** Same as **D** but for pairs of conditions differing in the 2nd (**E**) or 3rd (**F**) sequence position. **G)** Summary across arrays for decoders fit to listening activity generalizing to preparation and speech epochs. Binary LDA decoders fit as in B-D but on neural data joined over a 0.5s window of audio-cue listening. Decoders were evaluated on held-out trials for in-context listening windows as well as out-of-context preparation and speech windows of neural data. Average AUC over decoders shown. Error bars represent bootstraped 95% confidence intervals.

To test this, we fit decoders systematically across time points and tested their ability to predict sequence differences at past or future times spanning audio cue listening, preparation, and speech execution. Per-position decoders were fit to neural data from area i6v in T12 for 100ms time-bins during each trial epoch (same as Figure 1G) and then tested at all other trial time points. Decoders trained at one time point performed significantly above chance at many different time points (Fig. 4DEF). Surprisingly, decoders generalized not just between preparation and execution, but often across large parts of the full trial. For instance, decoders of 3rd-position phoneme identity fit during the cue period were significant throughout the audio cue, preparatory, and speech epochs (Fig. 4F, average 56% of time points in other trial epochs, 41% for 1st position, 63% for 2nd, 66% for 3rd; one-sample t-test of pairwise AUC against chance=0.5, *P*<0.05). To compare across microelectrode arrays, we quantified generalization by fitting decoders to a 0.5s window of the audio cue and assessing performance on the preparatory and speech epochs (Fig. 4C). In addition to T12 area i6v, decoders fit to T15 areas 55b and i6v also performed significantly better than chance when tested on the preparatory epoch (T12-area i6v: AUC=0.74 [0.72, 0.75]; T15-area i6v: 0.66 [0.62, 0.68]; T15-area 55b: 0.59 [0.56, 0.61] bootstrapped 95%CI, chance=0.5), and even better during the speech epoch (T12-area i6v: AUC=0.74 [0.71, 0.75]; T15-area i6v: 0.73 [0.70, 0.75]; T15-area 55b: 0.68 [0.65-0.69], bootstrapped 95%CI, chance=0.5, Fig. 4G, sliding-window version for T15 area i6v shown in Fig. S7). Generalization was not largely driven by a single position (Fig. S7D) and was not limited to training decoders on the audio-cue listening epoch (Fig. S8). In sum, phoneme sequence representations in precentral gyrus at least partially persisted between cue listening, preparation, and speech and do not exhibit strict orthogonality between preparatory and execution subspaces.

### Preparatory activity represents planned real words and is primarily limited to the 1st word in a sequence

Fluent speakers can effortlessly plan and produce long sequences of words. Psycholinguistic studies show that reaction times increase linearly with the number of words to be spoken, with each additional word adding roughly 10 ms of delay^2^ suggesting that word sequences are at least partially buffered before initiation of speech. To assess how sequences of words are represented in preparatory activity in motor cortex, participants were presented audio cues of sequences of three words in an instructed-delay task (Fig. 5A). Full factorial sequences of the three words [“clouds”, “vanish”, “abruptly”] were cued. In single electrodes, preparatory activity diverged primarily between conditions differing in the 1st word (Fig. 5B, 23/128 electrodes, Kruskal-Wallis H-test, FDR corrected, α=0.05). Preparatory activity on fewer electrodes were tuned to the 2nd word (4/128 electrodes), and none were tuned for the 3rd word, indicating that preparatory activity specific to the 1st word in the sequence was more broadly represented (note that 2^nd^ and 3^rd^ words become highly decodable later, during the production epoch of the trial).

**Figure 5:**
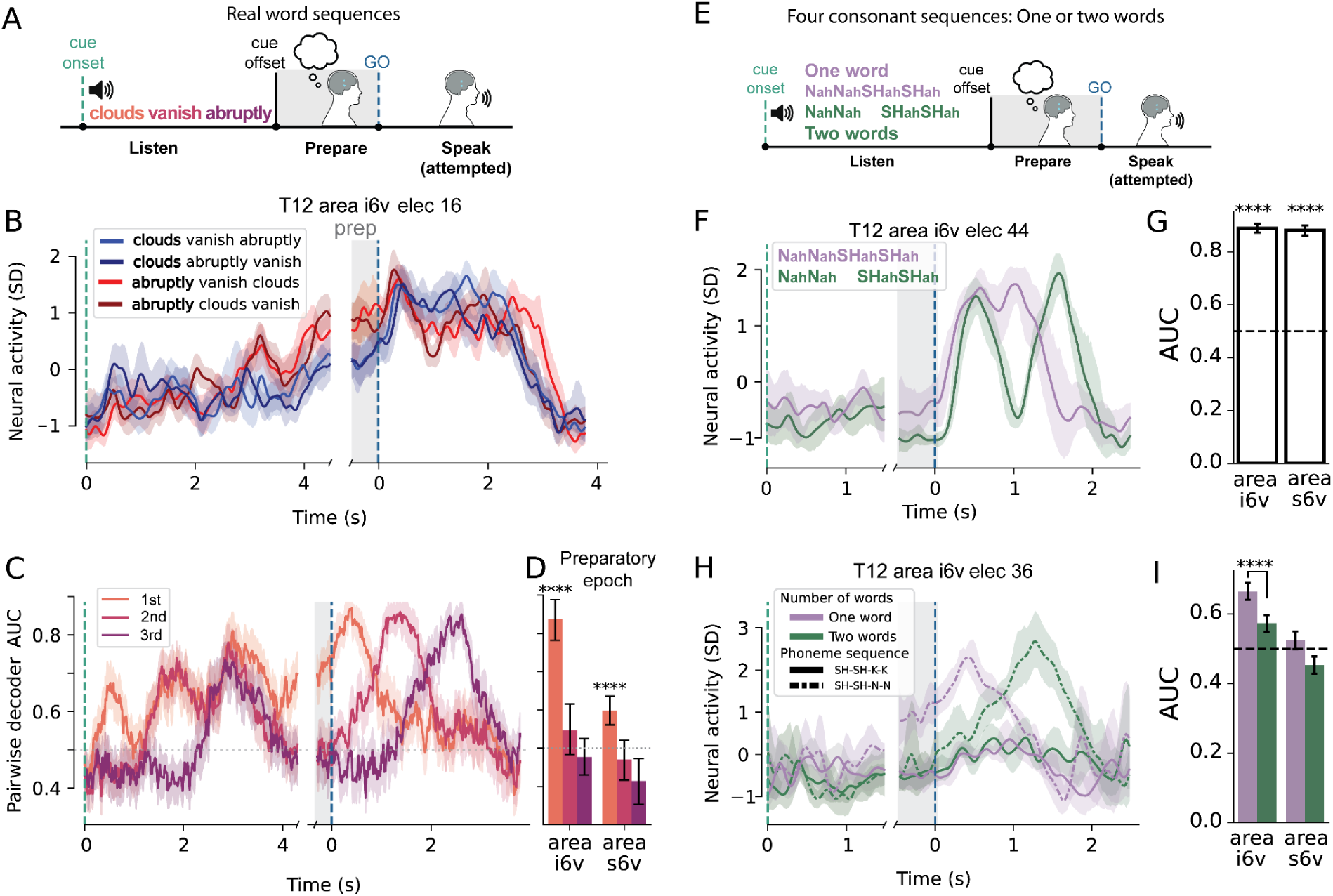
Preparatory activity tuning in area i6v is mostly limited to the 1st word in a sequence of words. **A)** All possible sequences of the words [“clouds”, “vanish”, “abruptly”] were presented in an audio-cued instructed delay task were cued. After a visuo-audio go cue, participant T12 attempted to speak the word sequence. Relative audio and go cue times are not to scale. **B)** Neural activity on a single electrode in area i6v in T12 for four conditions of real-word sequences. Gray shaded region indicates preparatory epoch — after all audio cues finished. Preparatory activity clusters by 1st word in the sequence. **C)** Population decoding from T12 area i6v for pairs that differ in a single position, analogous to Figure 1F (1st position example pair: “clouds vanish abruptly” vs “abruptly vanish abruptly”). Separability during the preparatory window is limited to the 1st word in the sequence. **D)** Summary of preparatory encoding over the preparatory epoch for areas i6v and s6v. Same analysis as in **C** for the average neural activity during the preparatory window. One-sample t-test versus chance=0.5, one-sided, Bonferroni correction for positions. **E)** Same task as in **A** but with a set of phoneme-balanced nonsense word sequences composed of four syllables that were either presented as one long word (purple) or two short words separated by a pause (green). A full factorial set of consonant sequences were cued. Relative audio and go cue times are not to scale. **F**) Neural activity on an example electrode in T12 area i6v that differed in preparatory tuning for the same phoneme sequence (N-N-K-K) presented as one long word (purple) versus two short words (green). **G)** Decoding between one vs two words, grouped across all phoneme sequences, from population activity during a preparatory window. One-sample t-test versus chance=0.5, one-sided. **H)** Neural activity on an example single electrode in T12 area i6v in response to four conditions that differ either in number of words (green vs purple) or phoneme sequence (solid vs dashed). The two-word conditions (green), which have the same first word (SHahSHah) have similar preparatory representations (shaded gray), but the corresponding one-word conditions (purple) do not. **I)** Population decoding between pairs of sequences that have the same starting phonemes but differ in the 3rd or 4th sequence position. In the two word context the differences are only in the second word. Decoding performance is significantly larger for one word conditions than two word conditions in area i6v, which performed better than chance. Two-sample, one-way t-test. (**BCFH**: Shaded intervals are bootstrapped 95% CI. Light gray shading indicates preparatory activity aligned to go-cue. PSTHs show smoothed, trial-averaged spike-band power. **DGI**: Error bars are bootstrapped 95% CI. \*\*\*\**P*<0.0001)

To probe these results at a population level, we fit pairwise decoders for 100ms sliding windows to sequences that differed in a single sequence position (e.g. 1st position: “abruptly clouds vanish” vs “vanish clouds vanish”). Aggregated results from all pairwise comparisons reveal that in T12 areas i6v and s6v, preparatory tuning was strongest for conditions differing in the 1st word (Fig 5CD, i6v: AUC=0.75 [0.67, 0.81], *P*=6.21E-07; s6v: 0.60 [0.56, 0.64], *P*=4.36E-05, bootstrapped 95% CI, one-sided t-test with Bonferroni correction for multiple positions) with markedly reduced decodability for the 2nd and 3rd positions (2nd: i6v: AUC=0.55 [0.48, 0.62], *P*=0.18; 2nd s6v: 0.47 [0.42, 0.52], *P*= 1.0; 3rd i6v: 0.48 [0.43, 0.53], *P*=1.0; 3^rd^ s6v: 0.41 [0.35, 0.47], *P*=1.0; bootstrapped 95% CI, one-sided t-test with Bonferroni correction for multiple positions, Fig. 5CD). These results suggest that rather than representing multiple words simultaneously, preparatory activity is primarily tuned for the 1st word.

To assess the relationship between phoneme sequence preparation and word boundaries, we instructed T12 to speak nonsense phoneme sequences of identical length and content, but either as a single word or pair of words. Audio cues consisted of four-syllable consonant sequences with fixed vowels (CVCVCVCV) and 3 possible consonants (K, N, SH) presented as either one four-syllable word (e.g., *KahNahSHahKah*) or two, disyllabic words (e.g., *KahNah SHahKah*, Fig. 5D). Thus, total phonemic content was matched across conditions, differing only in phonological word boundary^36^. Preparatory activity on single electrodes differed significantly between the one- and two-word contexts (92/256 electrodes, *P*<0.05, two-way t-test, FDR corrected, α=0.05, Fig. 5F). At the population level, decoders trained to classify the presence of a word boundary from preparatory activity performed well above chance (Fig. 5G; i6v: AUC=0.89 ± 0.02, *P*=6.3E-12; s6v: AUC=0.88 ± 0.02, *P*= 1.7E-11; 95% CI, one sample t-test over cross-validation folds). In other words, word boundaries significantly affected preparatory activity even for sequences with the same overall phoneme content.

We next asked how segmentation into two phonological words affected the structure of preparatory representations. From the results above, we hypothesized that two-word sequences with the same 1st word would be similarly represented in preparatory activity, but that the same phoneme sequences instructed as a single four-syllable word would be more distinct. This pattern was observed on an example electrode in area i6v in T12, where preparatory activity overlapped between two-word conditions with the same 1st word (*SHahSHah KahKah* vs *SHahSHah NahNah*; difference in average z-scored spike-band power Δ=0.231SD ± 0.451, *P*=0.622, standard error of the difference, Welch t-test), but the same phoneme sequences segmented as single words were significantly different on the same electrode (Fig 5H, focusing on preparatory period, denoted by shaded gray; *SHahSHahKahKah* vs *SHahSHahNahNah;* Δ=-2.059SD ± 0.356, standard error of the difference, *P*=0.00014). To assess the effect of word segmentation at the population level, decoders were fit to distinguish between pairs of phoneme sequences that only differed in the 3rd and 4th sequence position but were matched in the 1st and 2nd positions. Decoders were fit to the same pairs of conditions separately in the one-word and two-word context (see Methods for details). As expected from previous results, in the one-word context decoders performed significantly above chance (Fig. 5I; area i6v mean AUC: 0.665 ± 0.026, *P*=8.12e-30, 95% CI, one sample t-test) indicating that preparatory activity is tuned to the 3rd and 4th sequence position when preparing to speak a single, four-syllable word (Fig 5I). However, the same analysis for phoneme sequence in the two-word context performed significantly worse (ΔAUC = -0.09; *P*=2.9E-7, Fig. 5I). Therefore, the presence of a phonological word boundary resulted in weaker tuning for the 3rd and 4th sequence positions, in line with those positions now belonging to the second word in a two-word sequence. Together, these results indicate that preparatory activity in speech motor cortex is strongly affected by phonological word segmentation and has a single-word planning horizon.

## Discussion

The present results reveal that preparatory neural activity in human motor cortex contains a structured representation of long articulatory sequences. Phonemes across positions share subspaces with systematic geometric relationships, as opposed to independent orthogonal “slots” with a separate sequence-length encoding dimension. Phoneme representations generalize across listening, motor preparation, and attempted speech production epochs, suggesting a unified neural code linking perception, planning, and execution of spoken sequences. Finally, the limit of speech preparation is not determined by a fixed number of phonemes, but instead appears strongly affected by the 1st word boundary in a planned sequence of words. Other regions may be more strongly involved in preparing multi-word representations^24,37,38^.

### A gradient of preparatory representations across precentral gyrus

Despite their close anatomical proximity and, in some cases, shared functional parcellation (e.g. areas i6v and s6v), subregions of precentral gyrus exhibited markedly different preparatory tuning properties. Preparatory activity in area i6v and 55b encoded phoneme sequences spanning entire words, whereas nearby regions were primarily tuned to the initial articulation. Additionally, regions like area 4 while strongly tuned to the first phoneme did not contain decodable representations of higher-level features of planned words such as the length of the sequence. These findings indicate that preparatory representations are not uniformly distributed across motor cortex, but instead vary systematically across fine spatial scales. This is consistent with recent work demonstrating substantial differences in single-neuron tuning properties across precentral gyrus on the scale of millimeters^39^.

In the present study, preparatory tuning was largely limited to a single upcoming word. However, recent work has shown that parts of area 55b encode multi-word or sentence-level structure^24^. Taken together, these results suggest that preparatory representations in precentral gyrus may be organized along a functional gradient, with different subregions preferentially representing linguistic structure at distinct hierarchical levels^11,40,41^. Such an organization would support flexible speech planning through distributed, locally specialized representations rather than through a single centralized planning locus. Under this view, area i6v may function as a more downstream planning stage, converting higher-level sentence or phrase representations maintained in areas such as 55b into a concrete articulatory plan for the immediately upcoming word.

Despite longstanding theories implicating Broca’s area in the motor planning and sequencing of speech^11,40,41^, preparatory tuning was weak or absent in both microelectrode arrays targeting area 44 of inferior frontal gyrus. Although these data do not rule out speech-related representations in other subregions or at different spatial scales^42^, they are consistent with accumulating evidence that the contribution of area 44 to speech production diverges from classical articulatory motor-planning accounts^10,21,43^.

### Implications for intracortical speech BCIs

Intracortical speech BCIs have demonstrated the feasibility of restoring functional communication in individuals with paralysis^22^. Still, challenges remain in replicating such high performance in new participants, in part because the most informative cortical targets can be challenging to localize preoperatively. Consistent with this, we observed sharp differences in tuning between nearby array locations, underscoring the need for improved functional targeting. Regions previously associated with strong speech decoding performance^21–24,44^, also exhibited robust tuning during both speech preparation and auditory cue processing in the audio-cued instructed-delay paradigm. Together, these findings suggest that presurgical functional localization strategies could benefit from incorporating audio cue instructed delay tasks, which minimize movement-related artifacts during audio-cue listening while engaging similar neural representations as speech production. Such approaches may enable more precise, individualized targeting of highly informative regions such as areas i6v and 55b, as well as tailoring BCI algorithmic priors to the structure of encoding, improving the reliability of future intracortical speech BCI deployments.

Our findings on the geometry and temporal stability of speech preparation may have practical implications for the design of BCIs. We observed that phoneme representations persist with similar structure across cue processing, delay, and attempted execution—spanning timescales of several seconds. While modern BCIs use highly expressive architectures that are not tailored to a particular assumption regarding the underlying structure, such as recurrent neural networks (RNNs) and transformers, knowledge of the underlying signal structure may be relevant in cases where there is not enough data or bandwidth to use such architectures. For example, specialized RNN architectures like LSTMs were developed specifically to store and allow access to information that is present in the distant past but no longer available at the cost of added complexity in terms of states and parameters. This may not be a favorable tradeoff in some cases if our results on the stability of decoding generalize. On the other hand, our finding that encoding is compositional and close to linear suggests that decoding will benefit from nonlinear models as linear compositional encoding results in different conditions residing in tree-like structures that are not typically linearly separable.

### Simultaneous preparation of multiple speech sequence elements with context-dependent boundaries

Our work finds both commonalities with and substantial departures from previous studies of preparatory activity. In relation to primate reach studies — where Zimnik et al. (2021) found preparation of only the very next reach in premotor cortex — we found that for speech, preparatory activity in area i6v and 55b can be decoded to accurately determine not just the 1st phoneme in a sequence, but multiple phonemes deeper in the sequence. Xie et al. (2022) found encoding for multiple sequence elements in prefrontal cortex, but they were arranged in orthogonal subspaces for different positions in a sequence. In contrast, we find that positions share decoding subspaces. There are numerous possible explanations for these overall differences: the different brain areas recorded, the fact that these studies involve different species, extensive task-training on a fixed set of actions in model systems vs. natural speech, the difficulty of relating one type of movement, a reach, to another such as a saccade or the (attempted) articulation of a phoneme. Along these lines, Zimnik et al. make an insightful observation that in the study of voluntary movement, what counts as a single "voluntary movement", especially for sequences, is ill defined. Instead, they suggest discrete preparation as a way to define a voluntary movement. Relatedly, we found that the instruction to parse the same sequence into one or two words causes preparation to diminish at the word cutoff, i.e., the same phoneme position that was prepared before movement onset in the one-word context is not prepared in the two-word context. Characterizing more deeply the circumstances that change the window of preparation and whether they are consistent across different speech-motor areas may thus not only be important for empirical design of BCIs but may shed light on the nature of how speech is parsed by the brain into components, a subject of intense interest in the neurobiology of speech.

Orthogonal subspaces for different positions would provide a natural mechanism for encoding sequences of variable length and content, since each position could be read out independently. In that context, our finding that phoneme representations share subspaces across sequence positions is surprising. One interpretation would be that the difference is due to the nature and structure of speech. Sequential reaches, where each movement is planned and executed as a discrete unit, lend themselves well to orthogonal subspaces. Speech, however, imposes fundamentally different constraints. First, the articulatory plan for a word is inherently compositional: a word is not merely a sequence of independent phonemes but a coordinated gestural trajectory whose components interact through coarticulation, with the vocal tract configuration for one phoneme beginning to form during production of the preceding one, and are in the context of a hierarchically planned sequence of words, or at an even higher level, sequence of semantic concepts. Shared subspaces could potentially more naturally accommodate this interaction by representing all sequence elements simultaneously in overlapping dimensions, while reserving separate dimensions for alternative features such as sequence-length, as we observed in this work, or prosody. Second, the speed of speech (typically up to 10–15 phonemes per second) may make it impractical to sequentially rotate representations through orthogonal subspaces; instead, the brain may favor a scheme in which all upcoming phonemes are encoded concurrently, albeit with position-dependent scaling. Third, the brain’s encoding geometry may reflect the hierarchical structure of speech itself. Phonemes are not independent units — they are constituents of words which are constituents of sentences — and the fact that neural preparatory activity decreases with word segmentation suggests that the motor plan for an upcoming utterance is segmented according to this hierarchical organization. Representing sequential positions in overlapping rather than orthogonal subspaces may then mirror this compositional structure: because the phonemes of a word are planned together, their representations share neural subspace dimensions, reflecting their membership in a common higher-level unit. Indeed, the encoding we find of a word is directly constructed from the encodings of its constituent phonemes. The cost of this arrangement is first that the resulting representations are not linearly separable (i.e., a nonlinear readout is required) and second that it limits the length of sequence that can be encoded as at some point the scaling down of encoding for deeper positions will cause the signal to reach the level of noise. Because language can be decomposed into hierarchically organized sequences at multiple levels of abstraction (sequences of phonemes, words, phrases, or semantic concepts), compositional sequence planning with a shared subspace for sequence elements and orthogonal dimensions for related features may represent a general neural strategy, flexibly recruited across cortical areas supporting both speech and higher-level language processing.

### Mirrored representation during audio cue listening, preparation, and attempted speech

Our results align with prior reports that regions of precentral gyrus are tuned during both speech perception and production^8,23,45–47^, and connect naturally to the mirror-neuron framework, in which individual neurons respond during both passive observation and execution of the same action^48^. Rather than focusing on single-site selectivity or meso-scale field potentials, we examined population-level representations and found that activity patterns were correlated across trial epochs: decoders generalized robustly between audio cue listening, the preparatory delay, and attempted speech. Such population-level mirroring provides a potential neural instantiation of the shared perceptual–motor codes posited by computational models of speech production, in which ventral premotor cortex links stored articulatory programs to their predicted auditory targets^12^. At first glance this sits in tension with ECoG evidence that ventral sensorimotor cortex responses during speech listening are organized along acoustic rather than articulatory features, and do not recapitulate the somatotopic pattern evoked by speaking the same syllables^49^. However, the two sets of findings are reconcilable: shared population subspaces for phoneme identity can emerge from heterogeneous single-electrode tuning, as decoders selectively pool across many electrodes, such that the dimensions carrying phoneme information end up aligning across epochs even when individual electrodes differ in whether they are more "auditory" or more "motor." This common low-dimensional code for phoneme identity can be obscured at the meso-scale which pools underlying activity in different unknown ways such that the spatial layout of articulator-specific tuning dominates.

An important caveat is that cue listening in our task was not purely passive in that it was the only source of information for the instructed articulatory sequence, so the mirrored perceptual activity we observe may reflect a mixture of auditory processing and active preparation. Nevertheless, other studies have demonstrated activation of ventral precentral gyrus during both passive listening and speech^23,45,50^.

While consistent at a high level with theoretical models of speech-motor programming in ventral premotor cortex through shared perceptual–motor codes ^12^, our finding that the same decoders generalize all the way from audio cue listening, through preparation to attempted speech were surprising. Indeed, the full extent of a sequence is not known until one hears the end of the sequence and thus even in a shared perceptual-motor area a reasonable hypothesis would have been that there would be some form of intermediate representation during the audio cue, that transforms once the precise sequence is known and potentially transformed again into a different preparatory representation. Had this been the case, decoders would have generalized poorly into later phases. From the point of view of BCI development, a combination of non-linear decoding (as we find), modest spiking rates and fast transformations across representations would have meant that very high channel counts may be necessary for faithful readout. Our results that suggest more stable representations may help explain the success of BCIs with more modest channel counts as well as guide future development.

### Limitations

Several limitations should be considered when interpreting these results. First, both participants had ALS and produced dysarthric speech, which may influence the temporal structure of motor preparation. The strong encoding of only the 1st word in multi-word sequences, for instance, could partly reflect the need for longer-than-average pauses between words due to dysarthria^51^. However, fluent speech in able-speaking individuals is still segmented into utterances, though sometimes not aligned with lexical word boundaries (e.g. “let me” → “lemme”), so while the boundary of preparation was at the word boundary here this phenomenon of segmentation of articulatory plans may generalize to able-speaking individuals, but at the level of phonological word boundaries. Future studies using imagined speech paradigms could directly test whether the word-boundary effect persists in a less articulatorily constrained paradigm. Similarly, exact annotation of vocalized articulation onset times cannot be precisely observed in this research design.

In summary, the findings presented reveal the neural substrate for an articulatory buffer in the ventral (area i6v) and middle (area 55b) precentral gyrus whose representational geometry supports both compositionality and flexibility in speech production for novel words. Future explorations of the anatomical localization and structure of preparation for whole sequences of words will continue to elucidate the neural substrate of hierarchical speech production and, in conjunction with these results, may contribute to speech BCIs as a neural prosthesis to restore communication.

## Acknowledgments

We thank participants T12 and T15 as well as their carepartners for their generously volunteered time and effort as part of the BrainGate2 pilot clinical trial; B. Davis, K. Tsou, S. Kosasih, M. Massood, B. Travers, and D. Rosler, for administrative support; This work was supported by an ALS Pilot Clinical Trial Award (AL220043) from the Office of the Assistant Secretary of Defense for Health Affairs; a New Innovator Award (NIH 1DP2DC021055) from the National Institutes of Health, managed by the National Institute on Deafness and Other Communication Disorders; a grant (872146SPI) from the Simons Collaboration for the Global Brain; a postdoctoral fellowship from the A.P. Giannini Foundation; support from the Office of Research and Development, Department of Veterans Affairs (nos. N2864C, A2295R, and A4820R); the Wu Tsai Neurosciences Institute; the Howard Hughes Medical Institute; Larry and Pamela Garlick; and NIDCD (nos. U01DC017844, R01DC014034); a graduate student fellowship from the Blavatnik Family Foundation; and the NSF GRFP. The content is solely the responsibility of the authors and does not necessarily represent the official views of the National Institutes of Health, or the Department of Veterans Affairs, or the United States Government.

## Methods

### Study participants

Data from two participants, referred to as T12 and T15 are reported in this study, both of whom gave informed consent and were enrolled in the BrainGate2 Neural Interface System pilot clinical trial (ClinicalTrials.gov Identifier: NCT00912041, registered June 3, 2009). Approval for this pilot clinical trial was granted under an Investigational Device Exemption (IDE) by the US Food and Drug Administration (Investigational Device Exemption #G090003), as well as the Institutional Review Boards of Stanford University (protocol #52060) and the University of California Davis (protocol #1843264). All research was performed in accordance with relevant guidelines and regulations.

Given the unique ethical and philosophical issues inherent to intracranial neural recordings in human participants, ethical oversight was integrated into the research program from its inception. All potential risks, including those associated with surgery, derive exclusively from the parent safety and feasibility clinical trials rather than from the present study; these trials were each reviewed, approved, and continuously monitored by local Institutional Review Boards. Participant protection was further strengthened through several overlapping oversight mechanisms, including ongoing review by local Clinical Oversight Committees with an independent medical monitor, a Data Safety and Monitoring Board, compliance with FDA requirements under our Investigational Device Exemption, oversight by NIH clinical trial offices, and active collaboration with the NIH BRAIN Initiative Neuroethics Working Group. Together, these measures ensured adherence to the highest standards of safety and ethical integrity.

All research activities were conducted in accordance with the Declaration of Helsinki, the Belmont Report, CIOMS guidelines, and the NIH BRAIN Neuroethics Principles. Within the parent trials, participant autonomy and privacy were supported through an extended, dialogue-based informed consent process that evolved over several months and accommodated changing communication needs. In addition, the deliberate integration of neuroethics expertise throughout the trials promoted ongoing evaluation of risk–benefit tradeoffs, participant well-being, and broader societal implications.

T12 was 68 years old at the time of data collection, left-hand dominant, with slowly progressive bulbar-onset ALS diagnosed at age 59 (ALS Functional Rating Scale–Revised [ALSFRS-R] score of 26 at enrollment). At the time of recording, T12 had been dysarthric for approximately eight years due to bulbar ALS. She retained limited voluntary use of her limbs and primarily communicated using a writing board or tablet device. During attempted speech, she was able to vocalize and produce some subjectively distinguishable vowel sounds; however, consonants produced in isolation were largely unintelligible, and neither consonants nor vowels could be reliably identified during fluent sentence-level speech attempts. In March 2022, four 64-channel silicon microelectrode arrays (1.5-mm electrode length) coated with sputtered iridium oxide (Blackrock Microsystems, Salt Lake City, UT) were placed in the left hemisphere. Array locations were selected using preoperative structural and functional MRI and individualized Human Connectome Project (HCP) cortical parcellation (see ^21^ for details). Two arrays were placed in HCP-defined area 6v of the ventral precentral gyrus (orofacial premotor cortex), and two in HCP-defined area 44 of the inferior frontal gyrus (part of Broca’s area). Data included in this study span post-implant days 181 to 1133.

T15 was 45 years old at the time of data collection, with ALS diagnosed at age 40. At enrollment, T15 had dysarthria and no functional use of his upper or lower extremities (ALSFRS-R score of 23). His attempted speech did not produce audibly distinguishable speech sounds and his breath control was more limited than T12. In July 2023, four 64-channel silicon microelectrode arrays (1.5-mm electrode length) coated with sputtered iridium oxide (Blackrock Microsystems, Salt Lake City, UT) were placed in the left hemisphere. Array placement targeting was guided by preoperative structural and functional MRI and individualized HCP cortical parcellation (see ^22^ for details). Two arrays were placed in HCP-defined area 6v of the ventral precentral gyrus (orofacial premotor cortex), one array in HCP-defined area 55b, and one array in HCP-defined primary motor cortex (area 4). Data included in this study span post-implant days 349 to 424.

CAUTION: Investigational Device. Limited by Federal Law to Investigational Use

### Presurgical functional MRI for surgical planning

Before surgical placement of arrays, participants completed structural and functional MRI scans to support presurgical planning. These scans were used to assess hemispheric lateralization of speech and language functions, guide surgical decision-making, and inform targeting of microelectrode array placement (see ^21,22,39^ for estimated array locations and additional details).

### Neural signal processing

Neural activity was recorded as voltage time series using a Neuroplex-E system (Blackrock Microsystems) and transmitted through a percutaneous connector via a tethered cable. Signals were analog band-pass filtered using a fourth-order Butterworth filter with cutoff frequencies of 0.3 Hz and 7.5 kHz, then digitized at a sampling rate of 30 kHz with 250 nV resolution. Digitized data were streamed to custom MATLAB- or Python-based software BRAND^52^ for subsequent digital filtering and feature extraction (details below).

To extract signals associated with neural ensemble activity^53^, voltage traces from each electrode were digitally high-pass filtered at 250 Hz using non-causal filtering, with a delay of either 1 ms (T15) or 4 ms (T12). Linear regression referencing (LRR)^54^ was then applied to reduce shared noise across electrodes.

Two complementary measures of neural ensemble activity were subsequently derived for each electrode using 20 ms time bins. Threshold-crossing counts were calculated as the number of times the high-pass filtered voltage exceeded a negative threshold set at −3.5 times the signal’s standard deviation. Spike band power was computed as the summed squared voltage within each bin. Both threshold-crossing count and spike band power provide estimates of local population spiking activity and have been shown to closely match sorted single-unit activity with respect to decoding accuracy and population-level structure^55–57^. To mitigate slow drifts in neural activity across the session^58,59^ the mean threshold-crossing count and mean spike band power for each electrode were subtracted from the corresponding samples within each block.

Electrode-specific thresholds and LRR filter coefficients were determined using data recorded from an initial diagnostic block at the beginning of each session consisting of an instructed delay task for attempted speech of a set of real and nonsense words.

For all analyses, per-electrode neural activity was z-scored relative to the whole-session mean and standard deviation unless otherwise specified.

### Data collection

All digital signal processing and feature extraction were performed on a dedicated acquisition computer. For T12 sessions conducted prior to December 2024, neural processing was implemented using Simulink Real-Time, while task control was handled in MATLAB using the Psychophysics Toolbox^60^. A separate Windows computer managed task initiation and termination and communicated with the Neuroplex-E system. Beginning in December 2024 for T12 sessions, and for all T15 sessions, both neural data processing and task execution were implemented using BRAND^52^, a modular, Python-based framework. Data was collected in a series of blocks, with the number of trials per block, rest between blocks, and total number of trials collected determined by participants. All research sessions were performed at the participant’s place of residence.

A complete summary of all data collection sessions is provided in Table S1.

**Table S1:** Summary of all data collection sessions. Differences between sessions reflect differences in participant abilities and comfort. For example, T15 requested go-periods be self-paced to allow time to regulate breathing between trials. Sequence type notation indicates sequence elements as consonant (C), vowel (V), or word (W); spaces indicate an auditory pause between words; subscripts indicate sequence elements that vary across conditions. All cue designs were full factorial and only t12.2022.10.25 did not include a do-nothing condition. Delay duration includes both audio cue presentation and a post-cue preparatory period.

| Session ID | t12.2022.09.27 | t15.2024.06.30 | t15.2024.09.13 | t12.2025.05.06 | t12.2022.10.25 |
| --- | --- | --- | --- | --- | --- |
| Cue type | Nonsense word | Nonsense word | Nonsense word | Real word | Nonsense word |
| Sequence Types | C<sub>1</sub>VC<sub>2</sub>VC<sub>3</sub><br>C<sub>1</sub>VC<sub>2</sub><br>C<sub>1</sub>V<br>VC<sub>1</sub> | C<sub>1</sub>VC<sub>2</sub>VC<sub>3</sub> | C<sub>1</sub>VC<sub>2</sub>VC<sub>3</sub><br>C<sub>1</sub>VC<sub>2</sub> | W<sub>1</sub> W<sub>2</sub> W<sub>3</sub> | C<sub>1</sub>VC<sub>2</sub>VC<sub>3</sub>VC<sub>4</sub>V<br>C<sub>1</sub>VC<sub>2</sub>V C<sub>3</sub>VC<sub>4</sub>V |
| Sequence elements | C: K, N, SH, T<br>V: AH | C: K, N, SH<br>V: AH | C: K, N, SH<br>V: AH | clouds,<br>vanish,<br>abruptly | C: K, N, SH<br>V: AH |
| Total conditions | 89 | 28 | 37 | 28 | 163 |
| Trials per condition | 18 | 21 | 15 | 18 | 7 |
| Delay duration (s) | 2.5 | 1.5 - 3.5 | 2-4.5 | 5 | 2.5 |
| Go duration (s) | 2 - 2.25 | Self-paced<br>(2 - 15) | Self-paced<br>(2 - 45) | 4 | 2.5 |
| Figure | Fig 1, 2, 3, 4,<br>S2, S3, S4, S5,<br>S6, S7, S8 | Fig 1, 2, 4, S2,<br>S7, S8 | Fig 3, S6, | Fig 5 | Fig 5 |

### Audio-cued Instructed-Delay Task to instructor sequences of articulatory gestures

To probe the neural representation of articulatory sequencing independent of lexical semantics, participants performed an instructed-delay nonsense word speech task. Stimuli consisted of sequences of alternating consonants and vowels with a fixed vowel (ARPABET: /AH/; IPA: /ʌ/) and a subset of articulatorily distinct consonants (T12: IPA: [/K/, /N/, /ʃ/, /T/], ARPABET: [/K/, /N/, /SH/, /T/]; T15: IPA: [/K/, /N/, /ʃ/], ARPABET: [/K/, /N/, /SH/]). Sequences were operationalized as consonant sequences embedded in a canonical CVCVC structure (e.g., three-consonant sequences: C₁V C₂V C₃).

Nonsense words were used to generate a phonotactically balanced stimulus set while minimizing lexical and semantic influences on speech planning. Reaction times for real and phonotactically matched nonsense words have been shown to be comparable, suggesting that articulatory sequencing and motor program assembly proceed largely independently of lexical status (S. & Wright C. E., Knoll R., Monsell S., 1980). A full-factorial design was constructed such that every consonant appeared in every sequence position, yielding all possible combinations of consonant identities across positions. In addition, a “do-nothing” control condition was included in which no audio cue was presented and participants were asked to remain still following the go cue.

Audio cues were presented as spoken utterances recorded by a native English speaker; no visual text cues were shown. Each trial began with audio cue presentation, followed by an instructed delay period of either fixed or uniformly randomized duration (Table S1). During cue presentation and the delay period, participants viewed a gray screen containing a centrally located red square. The go cue consisted of the square turning green accompanied by a brief auditory tone, after which participants attempted to reproduce the cued sequence. All sessions also included a “Do Nothing” cue in which no audio was presented during the listening trial epoch and the words “Do Nothing” was presented on the screen and participants knew to do nothing for that trial during the preparatory and speech epochs.

Participant T12 attempted overt vocalized speech, whereas participant T15 attempted mimed speech without vocalization due to inconsistent breath control associated with ALS. Participants occasionally closed their eyes for comfort; all task information was conveyed auditorily, and no visual stimulus content was required. Delay and go-cue timing parameters were adjusted as needed to accommodate participant comfort and individual speech rates. Participants reported that the consonant sequences were easy to perceive and could be reliably reproduced without errors. As all trial epoch transitions and cues were auditorily presented, participants often performed the task with their eyes closed as it reduced eye-strain.

The cue-listening epoch was defined as audio cue onset to audio cue offset. The preparatory epoch followed the end of all audio cues and was defined as a 0.8s window preceding the go cue. The attempted speech epoch was defined as a 1 second window centered on speech onset. Speech onset times were manually annotated using visual inspection of microphone recordings in both the time and time–frequency domains, using Audacity and Label Studio software. Detailed trial timing parameters, number of trials per condition, and consonant inventories are reported in Table S1.

### Single electrode tuning

Single-electrode selectivity was assessed using statistical tests applied to threshold-crossing counts for each electrode. The window over which threshold-crossing counts were summed varied and are described below. All statistical tests were evaluated at a significance threshold of α = 0.05, with Benjamini-Hochberg false discovery rate (FDR) correction for multiple comparisons where applicable.

#### Electrodes significantly different than rest

First, we assessed whether neural activity was modulated differently than rest on each channel. Threshold-crossing counts across all trial epochs were summed to get the total activity for that trial on each channel. For each CVCVC sequence cue and each recording channel, we compared the distribution of single-trial threshold-crossing counts for that cue to the distribution for a rest condition (Do Nothing). Welch’s unequal-variance t-test was applied for each cue-by-channel comparison. We applied FDR correction across all such tests and classified channels as significant at corrected *P* < 0.05. We then summarized the number of channels that showed a significant cue effect for at least one CVCVC cue.

#### Electrodes significantly tuned to 1st, 2nd, or 3rd position

To assess whether preparatory activity on single channels was tuned to the 1st, 2nd, or 3rd position consonant, trials were grouped by the consonant identity in a specific position (e.g. all trials with /K/ in the 1st position). For each channel the Kruskal-Wallis test (nonparametric one-way test for multiple independent groups) was applied to the per-trial activity distributions across consonant identity to determine whether phoneme identity at the position was separable. We applied false-discovery-rate (FDR) correction across all channel × position tests using Benjamini–Hochberg–style adjustment, and counted channels with corrected *P*<0.05 for each position. For nonsense-word instructed delay tasks with CVCVC sequences, only the 2nd and 3rd position were assessed in order to quantify tuning for future sequence elements beyond initial articulation. All channels across all 4 microelectrode arrays in both T12 and T15 were included.

A similar analysis was performed to assess per-position tuning in the real-word sequences task for the 1st, 2nd, and 3rd position.

### Peristimulus Time Histograms (PSTHs) (Fig. 1D, Fig. 5BFH)

Before computing PeriStimulus Time Histograms (PSTHs), neural features on single electrodes were smoothed with a one-dimensional Gaussian kernel (σ =60ms) over data from the entire recording session. Activity aligned to one of three task events (audio cue onset, go cue onset, or speech onset). The smoothed activity was then averaged across trials within each condition.

### Linear discriminant analysis decoding

Neural ensembles typically encode task variables as population-codes: distributed patterns whose condition-specific differences are often well captured by low-dimensional structure in high-dimensional neural activity space (e.g. 64 electrodes of a Utah array). Typically targeted dimensionality reduction is employed to define such a subspace. In particular, Linear discriminant analysis (LDA) is a supervised learning approach that finds directions in the activity space that separate out samples from different classes, in our case trials with different target sequences.

LDA has a closed form solution through the eigenvectors of the product of two matrices: the between-class scatter matrix **B**, which captures how far each class mean strays from the global mean, and the within-class scatter matrix **W**, which accumulates the covariance of samples around their own class mean. Let *K* be the number of classes (e.g. phonemes), *n_k_* the number of trials in class *k*, and 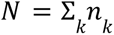 the total number of trials. From neural data **X** (n_trials × n_neural_features), with rows **x***_*i*_* and class labels *y* the between-class scatter matrix **B** and within-class scatter matrix **W** are estimated as:

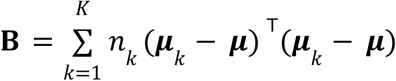

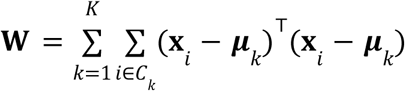

where *μ* is the global mean and ***μ*** is the class mean: 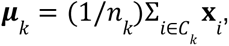, both row vectors over neural features for a single electrode. Each outer product is (n_neural_features × n_neural_features), accumulated over classes. *C_k_* denotes the set of trial indices for which the true class label is *k*.

In practice, the large number of parameters involved in the pooled within-class covariance matrix (number of features squared) leads to poor estimation. Instead, we regularized the estimation by replacing the pooled within-class covariance with its diagonal alone, i.e., treating features as conditionally independent given the class. Concretely, rather than inverting the full **W**, only its diagonal is retained: **D** = *diag*(**W**) (i.e. per-neural-features pooled within-class variances, with cross-neural-feature covariances set to zero). The optimal projection directions, or neural subspaces, that separate classes (i.e. the Fisher directions), are the eigenvectors of **D**^−1^**B** = λ**v**

For all LDA decoding analyses, neural activity features were per-electrode threshold-crossing counts and spike-band power for all electrodes on a single microelectrode array unless otherwise noted.

#### Binary LDA decoding between single-position difference pairs of conditions (Fig. 1EFG, Fig 4BCDE, Fig. 5CD)

Binary LDA classifiers were trained to discriminate between pairs of CVCVC sequences that differed in only a single consonant position (single-position difference pairs; example 1st position difference pair of conditions: **N**-AH-N-AH-SH, **K**-AH-N-AH-SH). Decoders were fit either to sliding 100 ms windows or to entire preparatory epochs. Decoder predictions were estimated using a discriminant score, calculated as the inner product between the trial’s neural activity projected into the LDA subspace and the difference between projected class means, evaluated relative to their midpoint. Thus the sign of the discriminant score indicates which class mean the trial was closer to. Classification accuracy was determined by the sign of the discriminant score on each held-out trial, while class probabilities for AUC were obtained by applying the logistic function to that score. Performance was evaluated on held-out trials using leave-one-out cross-validation Decoder performance was quantified on held out trials using classification accuracy when reporting individual condition pairs (Fig. 1E) and area under the receiver operating characteristic curve (AUC) when aggregating across multiple condition pairs (Fig. 1FG; Fig. 4B–E; Fig. 5C,D). For AUC analyses, predicted class probabilities were obtained for each held-out trial, and AUC was computed over all trials for each condition pair. Results were grouped by the consonant position in which the two sequences differed and averaged across pairs.

Ninety-five percent confidence intervals were estimated using bootstrap resampling of trials (10,000 iterations). Statistical significance of decoder performance was assessed using one-sample, one-sided t-tests against chance performance (0.5), with FDR correction applied across condition pairs, position, and arrays.

#### Cross-position decoder generalization (Fig. 2A)

To assess whether neural representations generalized across sequence positions, multiclass LDA decoders were trained to predict consonant identity at a single sequence position using all CVCVC trials (Fig. 2A). Decoders were evaluated using stratified 5-fold cross-validation. Performance was quantified using AUC, computed either in-context (evaluated on the same position used for training) or out-of-context (evaluated against consonants in other positions). Reported AUC values reflect averages across cross-validation folds. Confidence intervals were computed via bootstrap resampling of held-out test data (10,000 iterations), ensuring that no training data were included in the resampling procedure.

Because within-position decoding performance varies across arrays and positions, raw cross-position AUC values are difficult to interpret as measures of generalization — an array with weak within-position decoding may appear to generalize poorly simply due to a lower performance ceiling. To account for this, cross-position generalization was normalized relative to within-position decoding performance for each array and position. Specifically, normalized generalization was computed as the ratio of above-chance cross-position AUC to above-chance within-position AUC: (AUC_cross − 0.5) / (AUC_within − 0.5), such that a value of 1 indicates generalization equal to within-position performance and 0 indicates chance-level generalization.

### Neural geometry of shared encoding of 2nd and 3rd position consonants (Fig. 2BC)

To visualize shared neural structure between consonant positions, multiclass LDA models were trained to decode a subset of three consonants (N, SH, T) appearing in the 3rd sequence position to find an optimal two dimensional encoding subspace for the 3rd position (Fig. 2B,C). Conditions for the three classes were grouped by the 3rd position, ignoring 1st and 2nd position consonants: [*ah*ahN, *ah*ahSH, *ah*ahT] (* indicates a set of conditions with any consonant in that position). Models were fit using threshold-crossing counts from all electrodes and trained on 50% of the trials. The remaining trials were grouped by 3rd-position consonant identity, averaged, and projected into the two-dimensional discriminant subspace defined by the LDA model.

Class-specific displacement vectors for 3rd-position consonants were defined as vectors from the origin to the corresponding class means in discriminant space. The same held-out data were then regrouped by combinations of 2nd and 3rd position consonants and projected into the same subspace. 2nd-position displacement vectors were computed as the difference between the mean projection of a given 2nd-3rd sequence (e.g. *ahNahSH) and the corresponding 3rd-position-only baseline (e.g. *ah*ah*SH). A visual explanation is provided in Figure S3A-D.

#### Principal component analysis (PCA) for visualizing underlying neural geometry (Fig. S4)

While LDA identifies dimensions that maximally separate predefined classes, it does so by applying a transformation that can shear and rescale the underlying geometry to find the optimal linear subspace that separates target classes; PCA, by contrast, finds an orthonormal basis that captures the directions of maximum variance in the data without reference to class labels, thereby preserving the metric structure of the high-dimensional representational geometry up to a projection. To visualize the geometry of preparatory phoneme encoding across all sequence positions in area i6v we first computed the condition-averaged activity (threshold-crossing counts) for each electrode in a 0.7 second window before the go cue. Neural activity was summed within this window for each trial, then averaged across trials sharing the same condition to obtain 27 condition means (3 phonemes x 3 positions) in the full channel space. Principal component analysis (PCA) was applied to these 27 condition means, and the top 3 principal components (PCs) were retained for visualization. The condition-averaged variance explained from these 3 components was computed. All trials of these 27 conditions were used for PCA fitting and subsequently projected into the top 3 PCs for further analysis.

To assess whether 2nd and 3rd position encoding share a common subspace, we first averaged the 27 condition means by marginalizing over 1st-position identity, yielding 9 condition means (3 phonemes × [2nd, 3rd] position). A 2D plane was fit to these 9 points via singular value decomposition of the mean-centered matrix. This plane captured more than 90% of the variance among the 9 condition means, confirming that 2nd and 3rd position phoneme encoding lies in a shared 2D subspace.

We then constructed the tree-like structure hierarchically by computing displacement vectors from different marginalized averages over increasing sequence length: The grand mean served as the root; primary branches connected the grand mean to the 3rd-position phoneme means (thick lines, marginalized over 1st and 2nd position consonant); secondary branches connected each 3rd-position mean to the 2nd-position subdivisions (medium lines, marginalized over 2nd position consonant); tertiary branches extended to 1st-position subdivisions (thin lines, condition average of whole sequences). Branch color indicates phoneme identity at each level. The figure is shown from two viewpoints to better illustrate both the planar structure and the largely out-of-plane extension of 1st-position branches.

### Linear encoding model of phoneme sequences (Fig. 2D-G)

In light of the compositional geometry in Figure 2C, we modeled preparatory neural activity using a linear encoding framework that treats a planned phoneme sequence as the sum of position-specific phoneme contributions. The goal of this approach is to assess the similarity in neural representations of phonemes across sequence positions. Threshold-crossing counts across electrodes were summed over a 0.7 second preparatory window. Only significantly tuned electrodes were included (see Methods section “Electrodes significantly tuned to 1st, 2nd, or 3rd position” for more details) and threshold-crossing counts were z-scored per-electrode using the mean and standard deviation across all trials. Formally, z-scored threshold-crossing counts (*y*) were modeled as the sum of position-specific phoneme identity vectors, such that activity reflects the linear superposition of contributions from the first, second, and third position (**p***_i_* for position *i*):

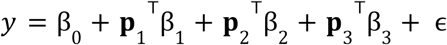

Model predictors (β) capture the tuning of each channel to phoneme identity at that position. An intercept term (β_0_ ) captures the grand mean activity across trials, and ɛ denotes residual variability not explained by the model. The predicted variable *y* contained threshold-crossing counts for electrodes significantly tuned to at least one sequence position from a single microelectrode array (0.7s preparatory window aligned to the go cue). The feature vectors [**p**_1_, **p**_2_, **p**_3_] are a one-hot encoding of phoneme identity at each sequence position(so each block **p***_i_* has dimension equal to the consonant inventory used in the task). Ridge regression was used (ɑ=0.5) as the dimensions of *y* was potentially large and the model was fit on noisy single-trials.

#### Model Performance

Model performance was evaluated using stratified cross-validation (3 splits), in which trials were grouped by condition to ensure balanced sampling of conditions across folds. For each fold, models were fit on ⅔ of trials and then predicted neural activity was computed for the held-out ⅓. Model predictions were pooled across all three splits and used to quantify performance using two metrics. First, single-trial R² was computed, reflecting the fraction of variance explained at the level of individual observations. Second, condition-averaged R² was computed using averaged responses for groups of like 2nd and 3rd position sequences (i.e. marginalizing over 1st position), providing a noise-normalized estimate analogous to a noise ceiling. For both R² metrics, multiple outputs were uniformly-averaged over meaning the variance explained of each electrode was weighted equally, regardless of the total variance of that electrode so as to not bias the metric towards prediction of a small number of high variance electrodes.

#### Cosine similarity summary

To analyze the structure of the learned encoding, model weights were refit on all trials without cross-validation. To assess the similarity of phoneme representations across sequence positions, we computed a cosine similarity matrix over coefficient vectors β_1_, β_2_, β_3_. Each predictor (i.e., each phoneme at each position) is associated with a weight vector across electrodes, and cosine similarity was computed pairwise across all such vectors (Fig. 2DE). To summarize the similarity across positions for each array with significant tuning for multiple sequence positions, we calculated the average cosine similarity between the same phoneme across different positions (e.g., phoneme /K/ at C1 vs. phoneme /K/ at C2, Fig 2F).

For both R² metrics and position-paired cosine similarity metrics, statistical significance was assessed via permutation testing (10,000 label shuffles), with *P*-values defined as the fraction of shuffled values exceeding the observed performance, and bootstrapped 95% confidence intervals were derived by resampling the pooled predictions with replacement 10,000 times and taking the 2.5th and 97.5th percentiles.

### Nonsense words of varying length (Figure 3)

Additional nonsense word sequences of varying syllabic length were constructed using the same consonant inventory (Fig. 3). Sequence lengths varied by participant due to session-specific trial yield constraints (T12: CV, VC, CVC, CVCVC; T15: CVC, CVCVC).

#### Sequence length and consonant decoding (Figure 3)

We first identified arrays with preparatory activity significantly tuned to sequence length. For each array, threshold-crossing counts and spike-band power were extracted over the preparatory window (0.6 seconds before the go cue, after all audio-cues have subsisted) and z-scored across trials. Sequence length was decoded using LDA (diagonal within-class scatter approximation, described above) with stratified 5-fold cross-validation. The number of sequence-lengths varied across participants due to differences in the number of conditions that could be collected in a single session. Therefore, performance was quantified as normalized balanced accuracy: (BA − chance) / (1 − chance), where chance = 1/K for K sequence-lengths. Bootstrap 95% confidence intervals were computed by resampling out-of-fold predictions (n=10,000). Statistical significance was assessed via a one-tailed permutation test (n=10,000) in which trial labels were shuffled and full cross-validation was repeated, with *P* value equal to the fraction of null draws ≥ observed. Only arrays showing significant sequence-length tuning after Bonferroni correction were included in subsequent compositionality analyses.

To estimate a neural encoding space for both phoneme sequence-length and content, LDA models were fit to predict combinations of sequence-length and phonemic content. While LDA is a supervised model, it does not assume any relationship between similar sequence-lengths or similar initial phonemes; in fact, the model should optimally separate all classes from one another, attempting to break compositionality in favor of larger mean separation if possible. LDA models were fit to jointly decode 1st position consonant identity and sequence length (CV, CVC, CVCVC), yielding nine target classes (Fig. 3) from spike band power and threshold-crossing counts averaged over a 0.6s preparatory window aligned to the go cue (after all audio cues finished). Models were trained on 50% of trials. Decoding performance on held-out trials was assessed via balanced accuracy. Held-out trials were grouped by target class, averaged, and projected into the top three discriminant components for visualization.

#### Model shared phoneme representation across different sequence lengths

The displacement-vector compositionality model was applied to preparatory neural activity projected into the LDA discriminant space for each array turned to sequence-length. The LDA model was fit to jointly decode 1st-position consonant identity and sequence-length (as described above), using 60% of trials for training (stratified by class). Held-out trials were projected into the resulting discriminant components for all subsequent analyses.

For each array, condition means were computed by averaging held-out trial projections within each sequence-length × 1st-position-consonant combination (target classes of the LDA model). Phoneme displacement vectors were then estimated as the difference between a given condition mean and the overall sequence-length mean, separately for each sequence-length (Fig. 3E). These displacement vectors capture the direction in discriminant-component space associated with a given phoneme, independent of sequence-length. Compositionality was evaluated using a leave-one-condition-out prediction scheme. For each held-out condition (defined by a sequence-length and 1st-position consonant), the predicted neural representation was constructed as the sum of (i) the phoneme displacement vector estimated from held out conditions (a different sequence-length, scaled to account for differences in displacement magnitude across sequence-lengths), and (ii) a sequence-length displacement vector capturing the mean difference between sequence-lengths. Predictions were averaged across both held-out sequence-lengths for T12 who was cued to speak 3 sequence-lengths.

Model performance was quantified as explained variance (R²) computed separately for condition-averaged data (condition-averaged R²) and for single trials (single-trial R²). Because the model outputs were projections into LDA discriminant components — which are ordered by discriminability and therefore capture unequal amounts of task-relevant variance — R² was computed as a single multivariate statistic weighted by the variance of each component rather than averaged uniformly across outputs. Bootstrap 95% confidence intervals were computed using 10,000 resamples of condition indices (for explainable R²) or trial indices (for single-trial R²). Statistical significance was assessed via permutation test (n=10,000), in which trial-level condition labels were shuffled prior to computing displacement vectors and predictions; the *P*-value was computed as (fraction of permuted R² values ≥ observed)

#### Cross-positional phoneme correlation structure across sequence lengths

As a complementary, decoder-free analysis to probe the relationship between sequence-length and phonemic content, we computed pairwise Pearson correlations between population activity vectors averaged over trials sharing the same sequence-length, and phoneme identity in a single sequence position. For each unique combination of sequence-length, sequence position, and phoneme identity, the mean preparatory activity vector was computed across spike-band power and threshold-crossing counts for all electrodes on a single microelectrode array (0.7s preparatory window aligned to the go cue). Pearson correlations were then computed between all pairs of such averaged population vectors, yielding a matrix whose rows and columns were organized by sequence-length, then position, then phoneme identity within each sequence-length (Figure S6, left).

To isolate phoneme-specific representational similarity from variance attributable to sequence-length, a mean-subtracted version of the correlation matrix was also computed. Prior to computing the within-group averages, the mean population vector for each sequence-length was subtracted from all trials belonging to that sequence-length, removing the shared sequence-length offset from the neural activity before computing pairwise correlations (i.e. marginalizing out sequence-length from the neural activity). This procedure allows the correlation structure to reflect phoneme identity and position independently of any global difference in preparatory state across sequence lengths (Figure S6, right).

### Temporal Persistence of Neural Representations (Fig. 4)

To characterize the temporal evolution of phoneme decoding performance across task epochs, binary LDA classifiers were fit to pairs of CVCVC trials differing in a single consonant position while holding all other positions fixed (see *Binary LDA decoding between single-position difference pairs*). Neural activity was extracted in 100 ms windows slid in 60 ms steps across the full trial period spanning cue listening, motor preparation, and attempted speech. Decoder performance at each time point was evaluated using leave-one-trial-out cross-validation and quantified as AUC computed from predicted class probabilities pooled across held-out trials. AUC values were averaged across all phoneme pairs and fixed-context comparisons for each sequence position to yield a single decoding time course per position per epoch.

#### Cross-time decoder generalization (Fig. 4BCD)

To assess whether phoneme representations generalized across time points within and between task epochs, decoders trained at each fit time point were evaluated at every other time point, yielding a fit-time × test-time generalization matrix. For each fit time point and epoch, a binary LDA classifier was trained and assessed using leave-one-trial-out cross-validation: for each fold, the held-out trial’s neural activity was evaluated at all valid time points across all epochs (cue, preparatory, speech). AUC was computed by pooling predicted class probabilities across held-out folds for each decoder × test time. Due to variability in audio cue and speech offset times, fit times were restricted to windows containing at least 900 trials across all conditions.

#### Cross-epoch decoder generalization using epoch-averaged activity (Fig. 4E)

To quantify whether phoneme sequence representations from audio-cue listening persisted into the preparatory and speech epochs across multielectrode arrays, a summary analysis was performed in which neural activity was averaged over fixed epoch-level windows rather than assessed at individual time points.

#### Peak time identification

For audio-cue listening epochs and speech epochs, the timing of when a phoneme position is heard or spoken varies. Therefore, the windows over which neural data was averaged varied by position. For each array and sequence position, peak decoding times were identified separately for the cue listening and attempted speech epochs. Binary LDA classifiers were fit on sliding windows. This followed the same pairwise decoding procedure described above except models were tested on neural activity from the same time window the model was fit on. AUC values were averaged as described above and the time point yielding the maximum mean AUC within each epoch was taken as the position-specific peak decoding time for that epoch. Note that this within-epoch decoding sweep is distinct from the cross-time generalization heatmap described above, in that the classifier is both trained and tested at the same time point; it serves solely to identify the time of peak phoneme discriminability within each epoch as an anchor for the subsequent cross-epoch generalization analysis.

#### Cross-epoch generalization

Three fixed analysis windows were constructed per position: a cue window (0.5 s centered on the cue-epoch peak decoding time), a speech window (0.5 s centered on the speech-epoch peak decoding time), and a preparatory window (the 0.5 s interval immediately preceding the go cue). Neural activity within each window was averaged over time to yield three feature vectors per trial — one per epoch. Binary LDA classifiers were trained on cue-window feature vectors using leave-one-trial-out cross-validation. On each fold, the model was evaluated on the held-out trial’s feature vector from all three epochs, yielding within-epoch (cue→cue) and cross-epoch (cue→preparatory, cue→speech) generalization performance. AUC was computed from predicted class probabilities pooled across held-out trials and averaged across all phoneme pairs and fixed-context comparisons. The analysis was conducted on CVCVC trials only and applied independently to each array combination (Figure 4E). The same analysis, but with training and testing on all combinations of trial-epochs was performed as well for Figure S8.

### Real word sequence decoding (Fig. 5A–D)

#### Real-word Sequences Instructed Delay Task

A similar task design was used to cue sequences of real words. Full factorial sequences of three real words ([clouds, vanish, abruptly]) were constructed similarly to nonsense words to create a set of 27 conditions consisting of all possible combinations. These words were selected as many combinations form grammatically valid clauses and each word is phonologically distinct. Audio cues were constructed by concatenating recordings of individual words separated by fixed silent intervals (0.2s).

#### Single electrode tuning and population decoding

Single-electrode preparatory tuning was assessed using a Kruskal-Wallis H-test applied separately for each sequence position (1st, 2nd, 3rd word) and each electrode, using threshold-crossing counts summed over a 0.5 s preparatory window aligned to the go cue. *P*-values were corrected for multiple comparisons across electrodes using FDR (α = 0.05).

To assess per-position decodability over the course of the trial, binary LDA decoders were fit to pairs of sequences differing in a single word position while holding all other positions fixed, using 100 ms sliding windows across audio cue listening, preparation, and attempted speech epochs, as described above (see *Binary LDA decoding between single-position difference pairs*, Fig. 5C). To summarize preparatory tuning, decoders were additionally fit to neural activity averaged over a 0.5 s window immediately preceding the go cue using leave-one-out cross-validation (spike-band power and threshold-crossing counts, Fig. 5D). AUC values were averaged across all pairs for each position. Statistical significance was assessed using a one-sided one-sample t-test of mean per-position AUC against chance (0.5), with FDR correction across the three sequence positions and microelectrode arrays.

### Nonsense word sequences with phonological word boundaries (Fig. 5E–I)

To assess the influence of phonological word boundaries on preparatory representations, four-consonant sequences were constructed from a three-consonant inventory ([K, N, SH]) with a fixed vowel (AH), yielding 3⁴ = 81 unique CVCVCVCV phoneme sequences. Each sequence was presented in two conditions: as a single four-syllable word (e.g. KahNahSHahKah) and as two disyllabic words separated by a pause (e.g. KahNah SHahKah), for a total of 162 conditions. The phonemic content was thus identical across the one- and two-word conditions for each sequence, differing only in the presence of a phonological word boundary after the second syllable.

#### Single-electrode word boundary tuning (Fig. 5F)

To assess whether individual electrodes distinguished one-from two-word contexts during preparation, a two-sample Welch’s t-test was applied to threshold crossing counts summed over the 0.5 s preparatory window, separately for each electrode, comparing trials from the one-word versus two-word conditions collapsed across all phoneme sequences. *P*-values were FDR corrected (α = 0.05) across all electrodes and arrays.

#### Grouped word boundary decoding (Fig. 5G)

To assess whether population activity during the preparatory window could discriminate one-from two-word contexts regardless of phoneme sequence identity, an LDA model was trained to decode the number of words (one vs. two) from neural activity (threshold-crossing counts and spike-band power) averaged over the 0.5 s preparatory window, using 10-fold cross-validation, stratified by phoneme sequence to ensure that each fold contained a balanced representation of sequences. Performance was quantified as AUC averaged across folds, with 95% confidence intervals computed assuming performance across folds is normally distributed (± 1.96 × standard deviation over folds). Statistical significance was assessed using a one-sided one-sample t-test against chance (AUC = 0.5).

#### Effect of word boundary on preparatory representations of later sequence positions (Fig. 5H–I)

To assess how the presence of a phonological word boundary altered preparatory representations of later sequence elements, we compared conditions that shared identical consonants in the first two positions (C_1_, C_2_) but differed in the third and/or fourth position (C_3_, C_4_). First, peri-stimulus time histograms were generated for one such pair, either in the two-word context (*SHahSHah KahKah* vs *SHahSHah NahNah*) or the one word context (*SHahSHahKahKah* vs *SHahSHahNahNah).* Spike-band power was averaged over a 0.5 second preparatory window on a single example electrode 44 in in T12 area i6v. For each context, we tested the between-condition difference with a two-sided Welch’s t-test. We summarized effect size as the difference in condition means (Δ) and quantified uncertainty with the propagated standard error of the difference over trials:

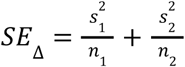

where *s_i_* and *n_i_* are the sample standard deviation and number of trials per condition *i*. Results are reported as Δ±*SE*_Δ_

#### Population decoding

Binary decoders were trained on pairs of conditions that shared identical consonants in the first two positions (C_1_, C_2_) but differed in the third and/or fourth position (C_3_, C_4_). All such pairs were identified exhaustively from the full factorial condition set. Binary LDA decoders were fit separately within the one-word and two-word contexts: in the one-word context, differing positions belonged to the latter half of a single four-syllable word (C_1_VC_2_V**C_3_**V**C_4_**V), whereas in the two-word context, the shared positions constituted the entire first word and the differing positions constituted the second word (C_1_VC_2_V **C_3_**V**C_4_**V). For each pair and context, a binary LDA classifier was trained on threshold crossing counts and spike-band power averaged over the 0.5 s preparatory window using leave-one-out cross-validation, with performance quantified as AUC computed from predicted class probabilities. Results were averaged across all qualifying pairs separately for the one- and two-word contexts, per cortical array. Statistical significance for each context was assessed using a one-sided one-sample t-test against chance; If at least one context performed significantly better than the difference in AUC between one- and two-word contexts was assessed using a one-sided two-sample t-test.

## Supplement

**Figure S1:**
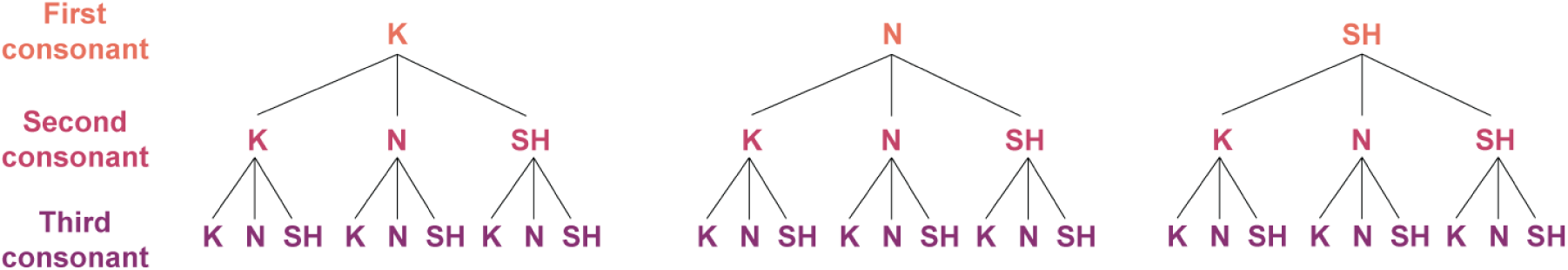
Schematic of full factorial cue design from a subset of consonants. Construction of a full-factorial cue set for 3 consonants (K, N, SH) in 3 positions creates a total of 27 total conditions. Related to Figure 1.

**Figure S2:**
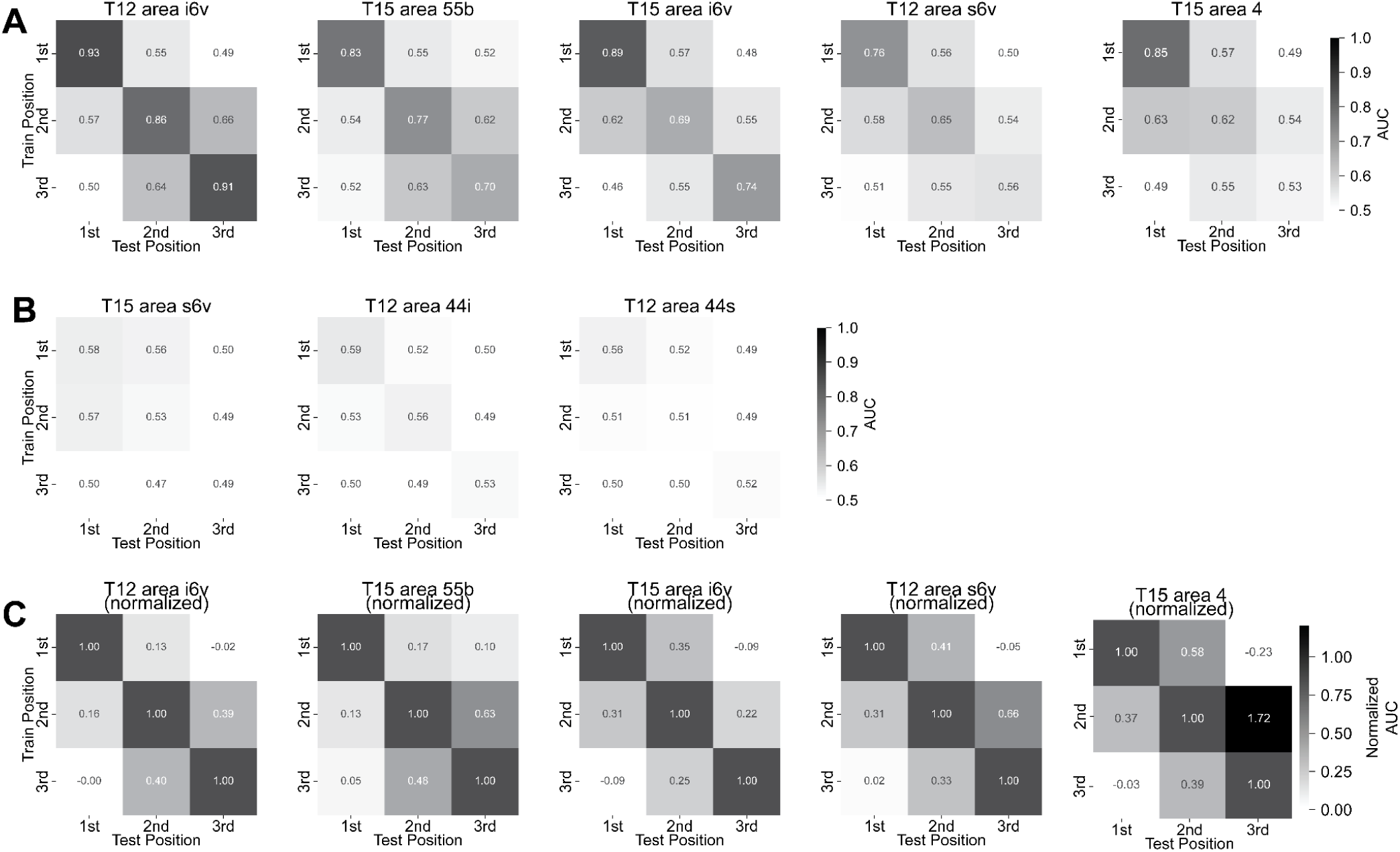
Cross-position generalization of phoneme decoding across all recorded arrays (related to Figure 2). **A)** For each microelectrode array, multiclass LDA decoders were trained to predict the consonant identity at each sequence position from preparatory neural activity and tested on held-out data labeled by all other positions, quantifying the degree to which phoneme representations generalize across positions (same analysis as Figure 2A; T12 area i6v reproduced here for comparison). **B)** Same analysis as panel **A** for arrays with weaker or absent preparatory tuning to phoneme sequences. **C)** Because within-position decoding performance can vary across positions, raw cross-position AUC values can obscure how well representations generalize relative to the performance ceiling of each position (e.g., in T15 area 4, 1st-position AUC is substantially higher than 2nd-position AUC). To account for this, cross-position AUC values were normalized by within-position performance (column-wise), using the ratio of above-chance cross-position to above-chance within-position AUC: (AUC_cross_ − 0.5) / (AUC_within_ − 0.5), such that a value of 1 indicates generalization equal to within-position performance and 0 indicates chance-level generalization. After normalization, generalization between the 2nd and 3rd positions is consistently stronger than between the 1st and 2nd positions in both T12 area i6v and T15 area 55b, consistent with the pattern of cosine similarities reported in Figure 2F.

**Figure S3:**
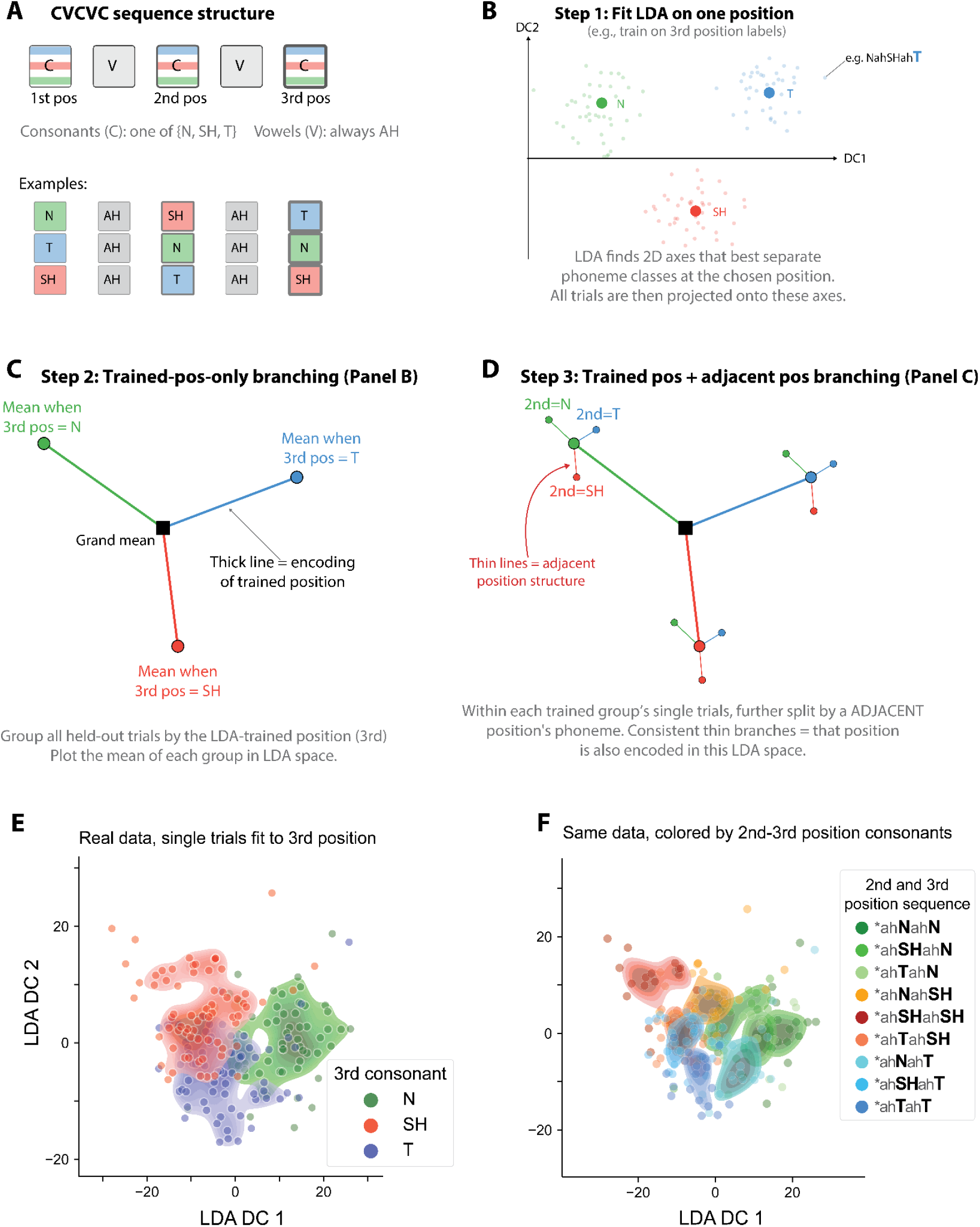
Explanation of the displacement vector analysis in LDA discriminant space (related to Figure 2BC). **A)** Schematic illustrating how full-factorial CVCVC sequences are operationalized as three consonant positions drawn from a three-consonant inventory. Vowels were fixed across all conditions. **B)** An LDA model is fit to decode a single sequence position (here, the 3rd) from preparatory neural activity, yielding two discriminant components (DCs) that maximally separate the three consonant classes. Simulated held-out trials projected into this DC space are shown for illustration: individual trials (small markers) cluster by 3rd-position consonant identity and class means (large markers) are well separated. Trials span many conditions (e.g., NahSHahT) but are colored solely by 3rd-position consonant (e.g., /T/). **C)** Held-out trials are grouped and averaged by 3rd-position consonant identity. A displacement vector is computed for each class as the vector from the grand mean of all held-out trials to each class mean (thick lines), representing the direction in DC space associated with each 3rd-position consonant. **D)** To visualize the encoding of adjacent positions, displacement vectors for the 2nd position are computed. Trials are now grouped and averaged by both 2nd- and 3rd-position consonant identity, yielding nine group means. For each 3rd-position class mean (derived in **C**), a 2nd-position displacement vector is computed as the vector pointing from that 3rd-position mean to the corresponding 2nd–3rd joint mean (thin lines), isolating the contribution of 2nd-position identity independent of 3rd-position context. Consistent orientation of 2nd-position displacement vectors for the same consonant across 3rd-position contexts (e.g., all thin red lines pointing in a similar direction for 2nd-position /SH/) and alignment with the corresponding 3rd-position displacement vector (thick red line) indicates a shared neural representation of the same consonant across positions. Note: panels **B–D** depict simulated data for illustrative purposes only; corresponding real data are shown in panels **E–F** and Figure 2B–C. **E)** Real held-out single trials from T12 area i6v underlying Figure 2B–C, projected into the LDA DC space and colored by 3rd-position consonant identity. Each cluster appears multimodal rather than unimodal. Threshold crossing counts were summed over the 0.7s preparatory window preceding the go cue, following audio cue offset. **F)** Same data as **E**, now colored by joint 2nd- and 3rd-position consonant identity. The multimodality visible in **E** is partially explained by sub-clustering according to 2nd-position consonant identity, consistent with the shared representational structure described in Figure 2.

**Figure S4:**
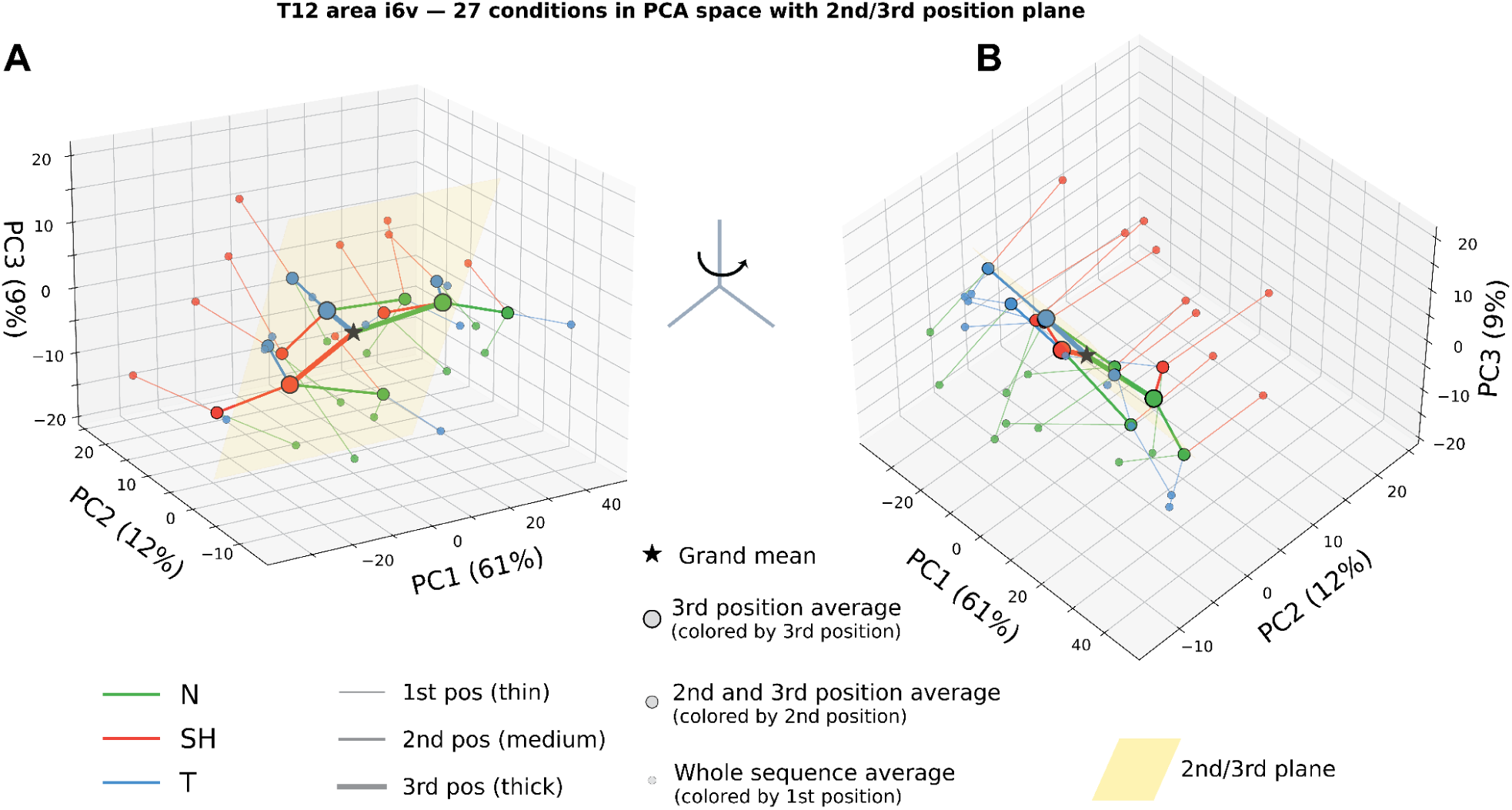
PCA-derived geometry of CVCVC sequences. **A)** Principal component analysis (PCA) provides an unsupervised approach to dimensionality reduction (compared to LDA), which is guaranteed to preserve the geometry of the original high-dimensional neural activity. PCA was performed on average neural activity (threshold-crossing counts over a 0.7s preparatory window from area i6v in T12) of whole sequences such that principal components are optimized to explain variance between average preparatory representations of phoneme sequences. Consonants subselected from {N, SH, T} to match Figure 2 resulting in 3^3^=27 total conditions. The top three principal components (PCs) captured 81% of condition-mean variance and were retained for visualization. Displacement vectors for the 1st, 2nd, 3rd position are shown. See Fig. S3 for visual explanation of analysis, extended here to the 1st position. A similar structure arises, with a 2D plane capturing more than 90% of the variance of 2nd-3rd position sequences (yellow, similar to Figure 2BC). Within this plane, displacement vectors are consistently aligned for the 2nd and 3rd position (medium and thick lines). **B)** The same data as in **A** are shown from a rotated view orthogonal to the 2nd-3rd position plane. 1st phoneme representations are organized largely orthogonally to the 2nd-3rd position plane. Related to Figure 2.

**Figure S5:**
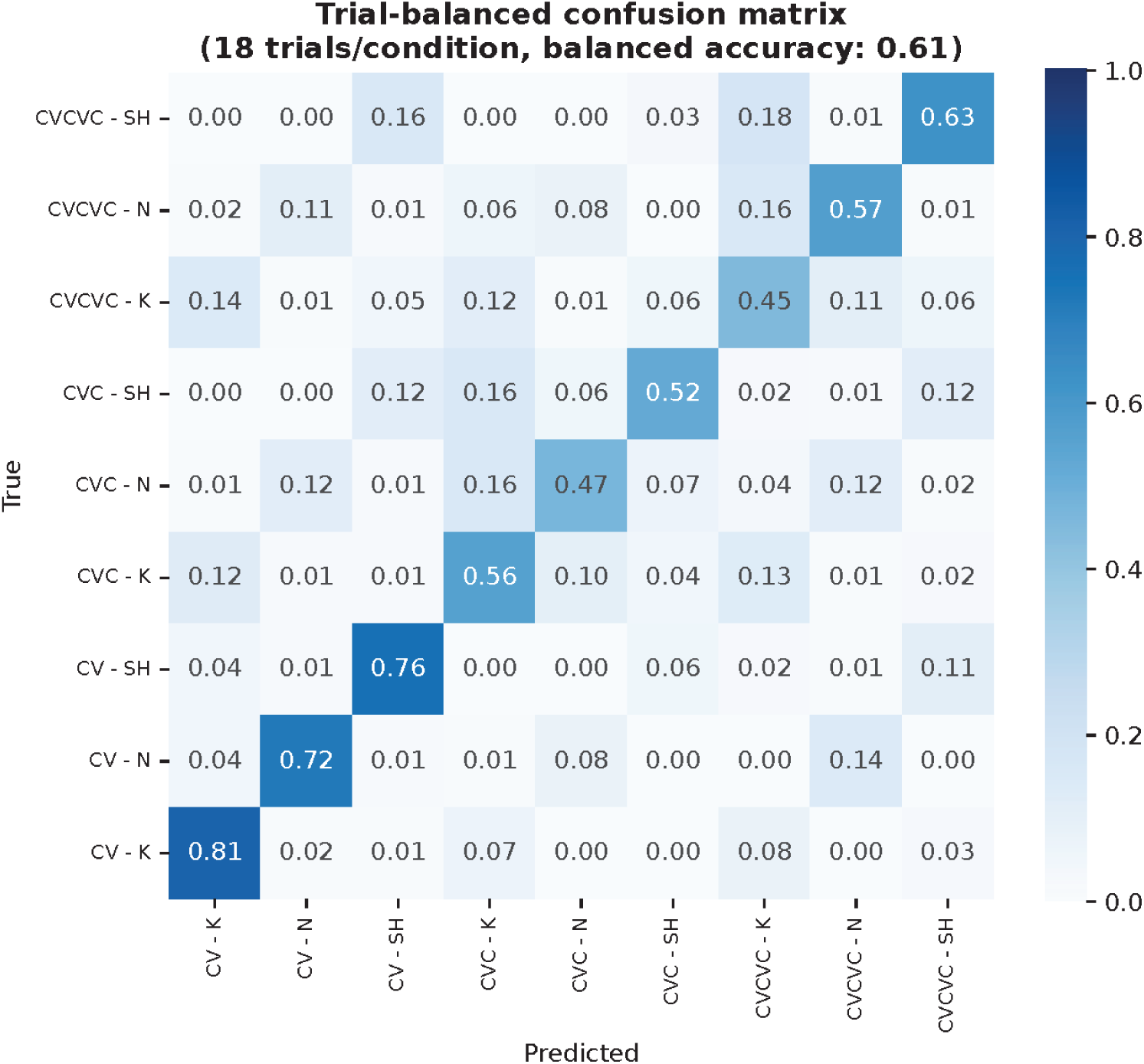
Trial-balanced decoding of sequence length and first-position phoneme identity from preparatory activity in T12 area i6v. In the factorial design used here, fixing trial counts per sequence condition results in fewer total trials for shorter sequence types, which contain fewer unique conditions (e.g., 3 CV conditions versus 27 CVCVC conditions for a 3-consonant inventory). The decoder in Figure 3B was therefore trained on all available trials, introducing class imbalance that may inflate apparent confusion among minority-class sequence lengths due to a difference in class probability. To assess whether the confusion pattern in Figure 3B reflects the underlying representational geometry rather than class imbalance, a trial-balanced decoder was fit after subsampling majority classes to match the trial count of the minority class. Decoding performance remained strong and broadly uniform across all sequence length × first-position phoneme classes, demonstrating that the confusion observed for shorter sequences in Figure 3B is due to trial imbalance.

**Figure S6:**
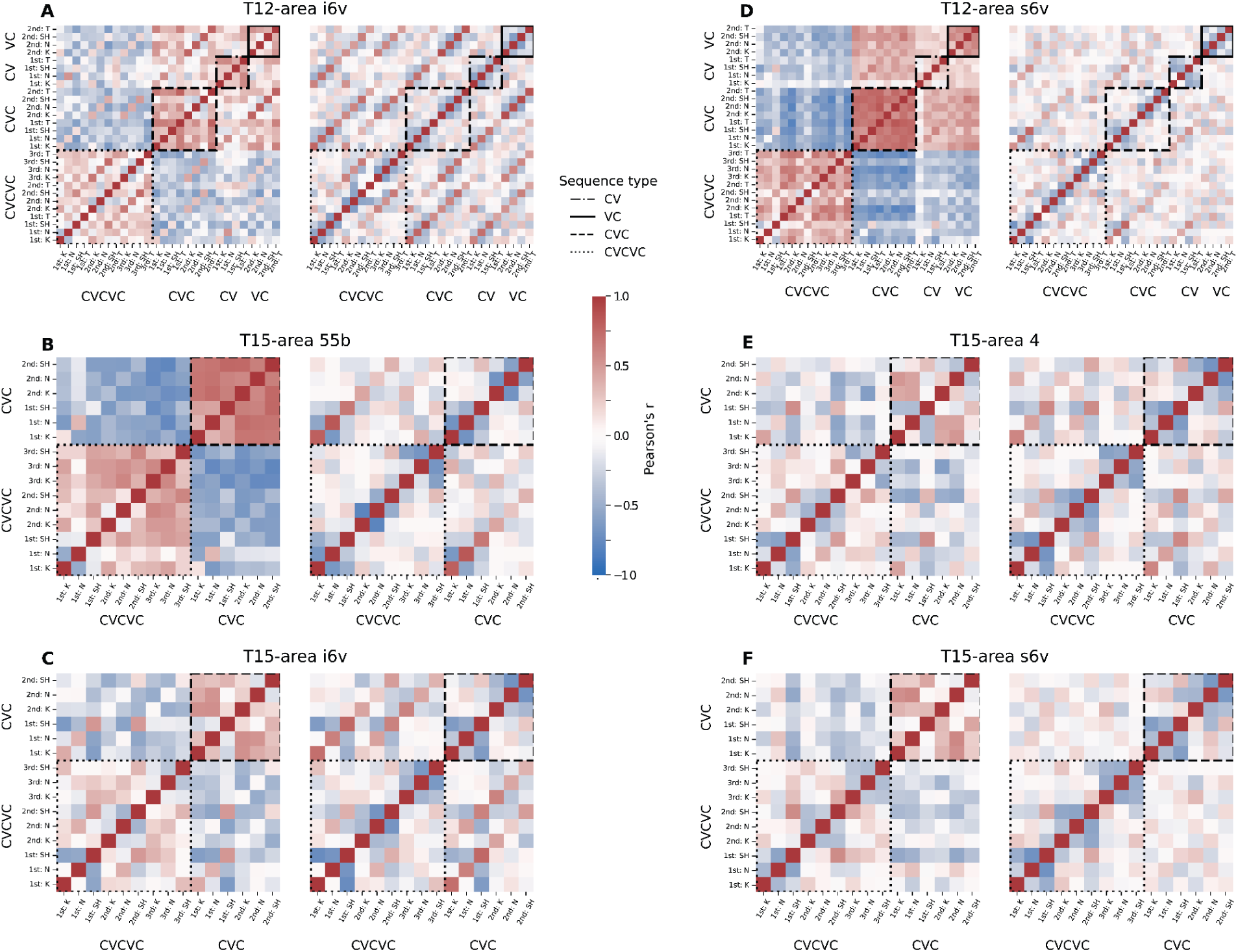
Like-sequence types are broadly correlated across phonemes and marginalizing out sequence-type reveals a shared phonetic code. Pairwise Pearson correlations between population activity vectors averaged over trials sharing the same sequence type, and phoneme identity in a specific sequence position. Each matrix entry reflects the correlation between two such averages — for example, the mean preparatory population state for the 2nd-position /K/ in CVC sequences versus the 1st-position /SH/ in CVCVC sequences — computed across spike-band power and threshold-crossing counts on all electrodes in a single microelectrode array during the preparatory window (0.6s prior to go cue). Diagonal squares outlined rectangles indicate entries sharing the same sequence type. Each pair of matrices show results for a single microelectrode array; Left: Raw correlation matrix. Right: Correlation matrix after subtracting the sequence-type mean population vector from each trial prior to averaging, isolating phoneme- and position-specific representational geometry from sequence-type-level offsets. Related to Figure 3.

**Figure S7:**
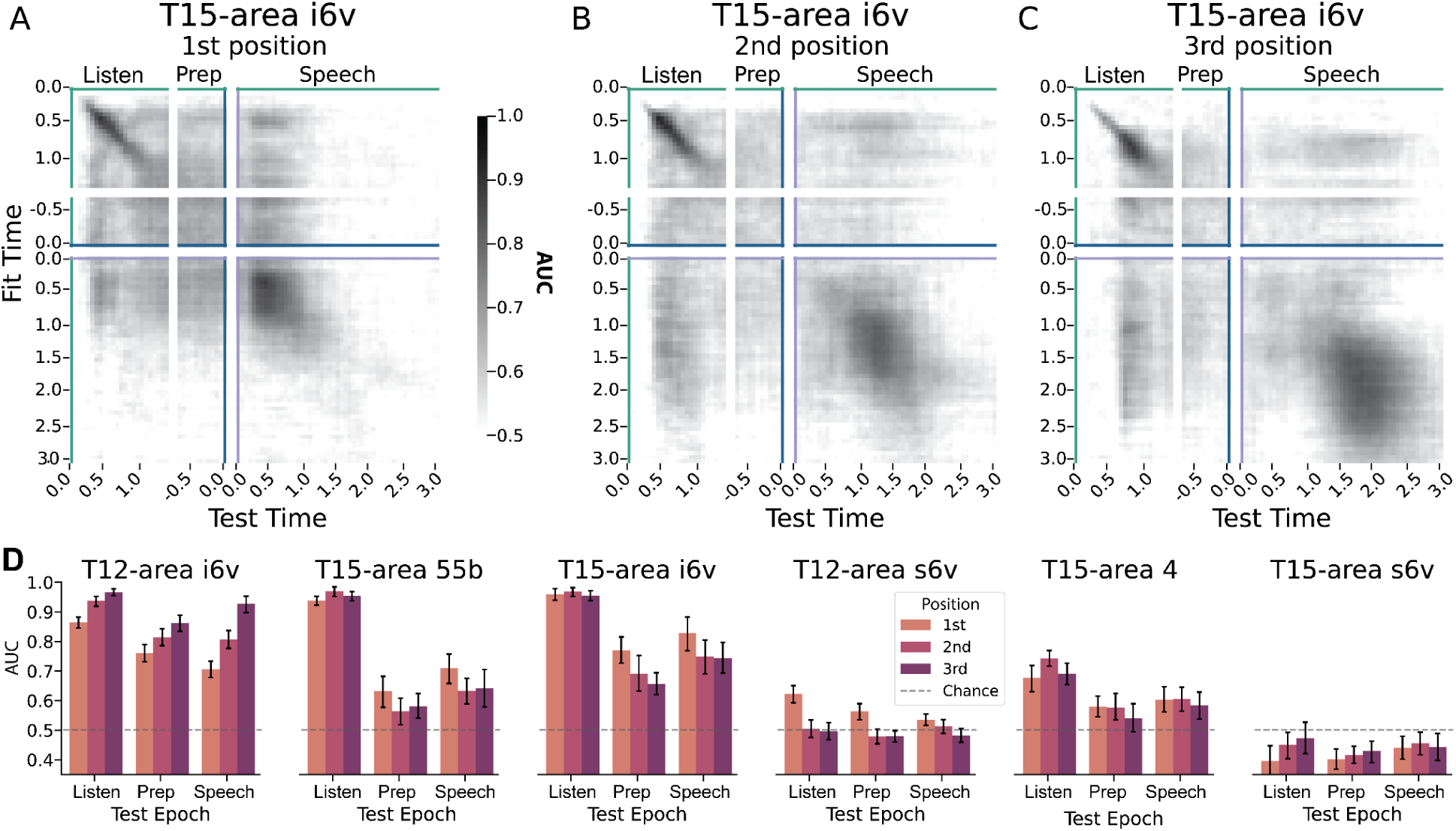
Cross-time generalization for T15 and per-position epoch generalization. **A)** Cross-time decoder generalization analysis of Figure 4D reproduced for T15-area i6v (1st position decoding performance). Decoders generalize broadly from listening to preparation and speech. Results were similar for 2nd (**B**) and 3rd (**C**) position decoders. **D)** Same cross-epoch decoding results as Figure 4G: decoders fit to predict between pairs of conditions from neural activity averaged over a 0.5s window of audio-cue listening and tested during all trial-epochs. Results here are plotted by sequence position (results in Figure 4G are for average over position). Error bars are 95% confidence intervals over decoders.

**Figure S8:**
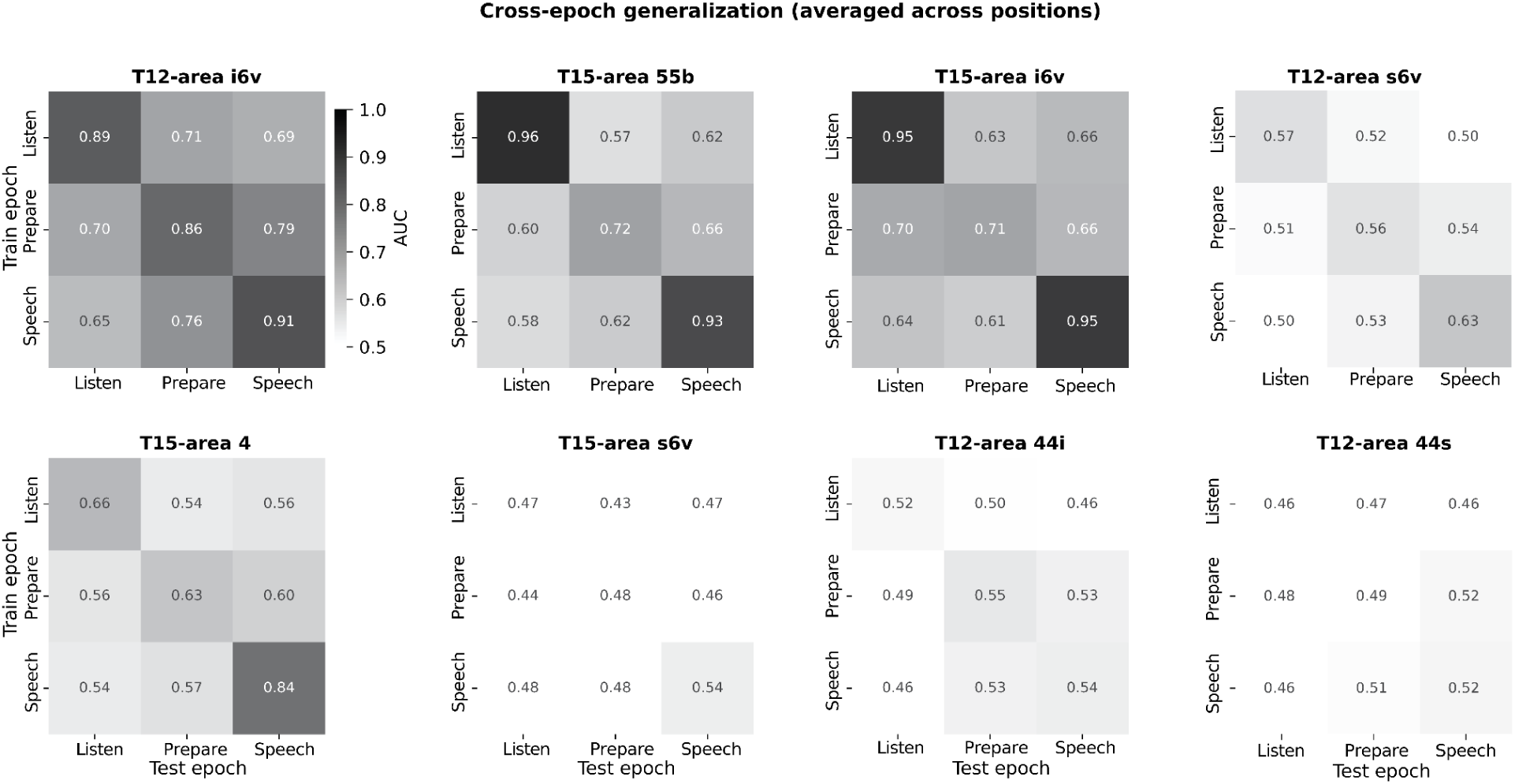
Cross-epoch generalization of phoneme decoding across all trial-epoch pairs, per array. Extension of Figure 4E in which decoders are trained and tested on all combinations of task epochs (Listen, Prepare, Speech), rather than training on Listen only. Each panel shows decoding performance (AUC, chance = 0.5) for a single array; rows indicate training epoch, columns indicate test epoch. Diagonal entries reflect within-epoch performance; off-diagonal entries reflect cross-epoch generalization. Values are averaged across sequence positions.s

